# Identifying the Biosynthetic Gene Cluster for Triacsins with an *N*-hydroxytriazene Moiety

**DOI:** 10.1101/495424

**Authors:** Frederick F. Twigg, Wenlong Cai, Wei Huang, Joyce Liu, Michio Sato, Tynan J. Perez, Jiaxin Geng, Moriel J. Dror, Ismael Montanez, Tate L. Tong, Hyunsu Lee, Wenjun Zhang

## Abstract

Triacsins are a family of natural products containing an N-hydroxytriazene moiety not found in any other known secondary metabolites. Though many studies have examined the biological activity of triacsins in lipid metabolism, the biosynthesis of triacsins has remained unknown. Here, we report the identification of the triacsin biosynthetic gene cluster in *Streptomyces aureofaciens* ATCC 31442. Bioinformatic analysis of the gene cluster led to the discovery of the tacrolimus producer *Streptomyces tsukubaensis* NRRL 18488 as a new triacsin producer. In addition to targeted gene disruption to identify necessary genes for triacsin production, stable isotope feeding was performed *in vivo* to advance the understanding of N-hydroxytriazene biosynthesis.

---

Triacsins are a class of acyl-CoA synthetase (ACS) inhibitors consisting of an 11-carbon alkyl chain and a terminal *N*-hydroxytriazene moiety (Figure 1). They were discovered almost four decades ago in *Streptomyces aureofaciens* ATCC 31442 through an antibiotic screening program and characterized as vasodilators.^[1–2]^ Being mimics of fatty acids, triacsins were the first ACS inhibitors to be discovered,^[3]^ and their activities in both bacterial and animal models have made them invaluable in the study of lipid metabolism.^[4–7]^ More recently, triacsins have been shown to have an antimalarial activity similar to artemisinin^[8]^ and an antiviral activity against rotaviruses.^[9]^ Though many studies have covered the biologic activity of the triacsins, the biosynthetic origin of this family of compounds remains elusive.

**Figure 1.**
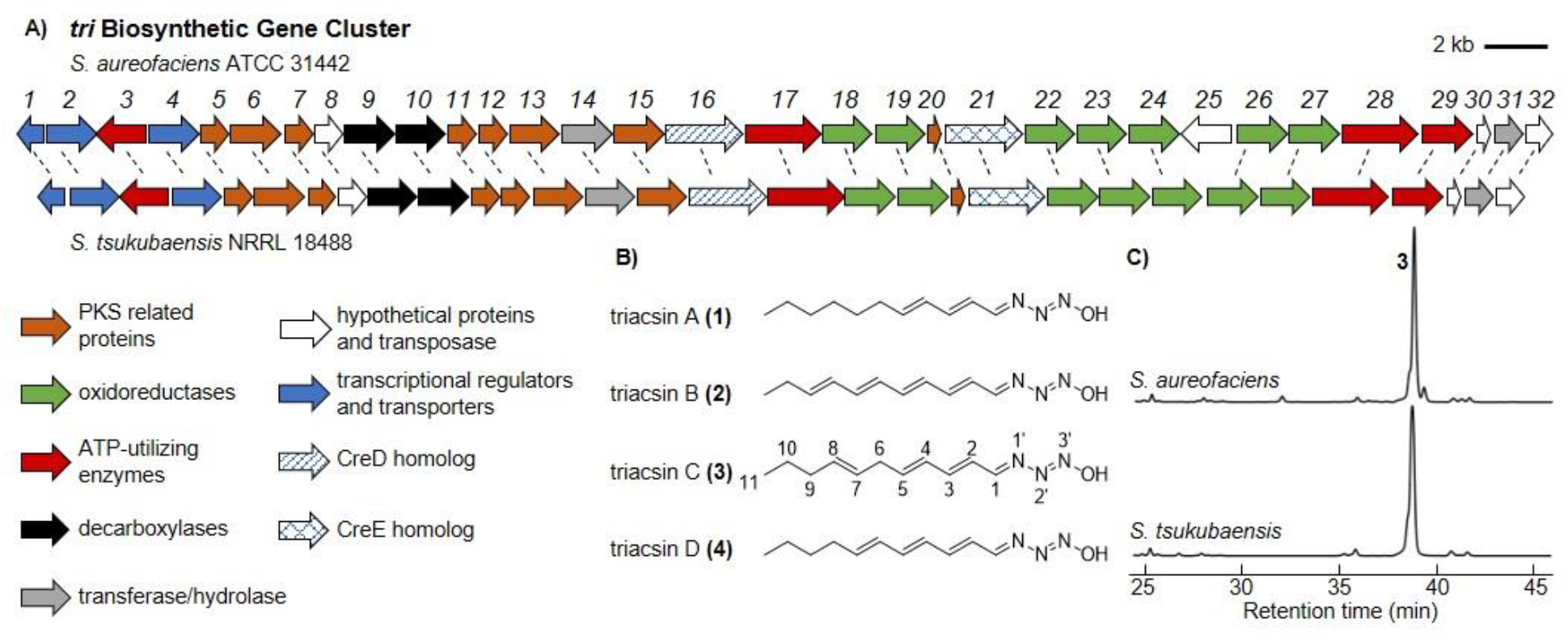
Identification of the triacsin *(tri)* BGC and a previously unrecognized natural producer. A) Schematic of the *tri* BGC as discovered in *S. aureofaciens* and the homologous BGC identified in *S. tsukubaensis.* B) All four of the originally reported triacsin congeners contain an 11-carbon alkyl chain and the unique *N*-hydroxytriazene moiety. C) UV traces (300 nm) confirming the production of 3 in *S. tsukubaensis.* In both strains, the congener 3 was produced as the major product and other congeners were not identifiable by UV in these traces.

Four triacsin congeners were originally identified which feature a conserved N-hydroxytriazene moiety and vary in the unsaturation pattern of their alkyl chains (Figure 1). The N-hydroxytriazene moiety, containing three heteroatom-heteroatom linkages, is rare in nature and has only been identified in the triacsin family of natural products. Given the prevalence of N-N and N-O linkages in synthetic drug libraries, there is a great interest in their enzyme catalyzed formation to complement the challenges in synthetic approaches.^[10–11]^ Up to date, multiple classes of enzymes have been characterized to catalyze N-O linkages such as cytochrome P450s and N-oxygenases which may be flavin-dependent or metal-containing.^[11]^ For N-N bond formation, few mechanisms have been examined until very recently. One chemical logic goes through the activation of one nitrogen to a more electrophilic species followed by intramolecular addition to a second nucleophilic nitrogen. For example, N5 of ornithine is hydroxylated by a flavin-dependent enzyme (KtzI) before the N-N bond formation promoted by a heme enzyme (KtzT) (Figure S1).^[12–13]^ Similarly, N6-hydroxylysine is conjugated to glycine followed by intramolecular nucleophilic attack from the glycine amino group catalyzed by a fusion protein (Spb40) consisting of a cupin and a methionyl-tRNA synthetase-like protein (Figure S1).^[14–15]^. An alternative mechanism for N-N bond formation features the reaction of an amine with nitrous acid that is generated through the sequential function of a flavin-dependent enzyme (CreE) on aspartate and a lyase (CreD) (Figure S1).^[16–20]^ For example, the diazo moiety found in cremeomycin is generated through the reaction of an aryl amine with nitrous acid promoted by an acyl-CoA ligase homolog (CreM) (Figure S1).^[21]^ Similarly, the phosphonohydrazide moiety of fosfazinomycin and the diazo moiety of kinamycin are both formed through the intermediacy of nitrous acid, although the dedicated N-N forming enzymes have not been identified in these cases (Figure S1).^[22]^ Compared to these N-N containing natural products, triacsins have three consecutive nitrogen atoms in the *N*-hydroxytriazene moiety, adding complexity to N-N bond biogenesis. Investigation of the biosynthesis of triacsins may determine if a known mechanism is used or if yet another chemical logic for N-N bond formation exists.

To identify the potential genetic context for triacsin biosynthesis, the genome of the known triacsin producer, *S. aureofaciens* ATCC 31442, was subjected to Illumina and PacBio sequencing, which resulted in 7.6 M nonredundant bases after assembly of paired sequence reads. The genome was annotated using RAST^[23]^ and biosynthetic gene clusters (BGCs) were predicted using antiSMASH.^[24]^ Genome mining for triacsin BGC candidates was performed by a local BLASTP analysis on the genome using probes such as polyketide synthases (PKSs) which may be involved in the alkyl chain formation, or enzymes from known N-N bond formation pathways such as KtzT, CreE and CreD. While no homologs were found for KtzT, our bioinformatic search turned up two putative BGCs encoding both CreE and CreD homologs (Figures 1 and S2). Cluster 1 spanned 30 kb and contained 21 open reading frames (ORFs). Cluster 2 spanned 39 kb and contained 32 ORFs. A targeted gene disruption deleting a 9 kb multigene region from cluster 1 did not affect triacsin production (Figures S3 and S4). In comparison, a similar multigene disruption in cluster 2 abolished the production of all triacsins (Figure S4). We then assign the cluster 2, designated *tri1-32,* as the putative triacsin BGC.

To further confirm the importance of this BGC in triacsin biosynthesis, we used an architecture search with the MultiGeneBlast tool^[25]^ looking for regions of homology in or surrounding the *tri* BGC within published genomes. Two additional microbes were identified containing BGCs highly homologous to the *tri* BGC (Figures 1 and S5); the marine actinomycete *Salinispora arenicola* CNS-205 (Accession #: PRJNA17109)^[26]^ and the well-studied tacrolimus producer *Streptomyces tsukubaensis* NRRL 18488 (Accession #: PRJNA389523).^[27–29]^ We then cultured these strains to examine their capability in producing triacsin family compounds or potential derivatives. While *S. arenicola* did not appear to produce triacsins in the tested conditions, triacsins were confirmed to be produced by *S. tsukubaensis* through UV and liquid chromatography-high resolution mass spectrometry (LC-HRMS) analyses (Figures 1 and S6). Similar to *S. aureofaciens, S. tsukubaensis* produced all four known triacsin congeners with triacsin C (3) being the major metabolite. Careful analysis of *S. tsukubaensis* cultures identified four additional compounds with UV spectra similar to 3 or triacsin D (4) and the same molecular formulas of C_11H17_N_3_O_2_ indicated by HRMS, suggesting that they were hydroxylated triacsin derivatives (5-8) (Figure S7). These results revealed for the first time that *S. tsukubaensis* is a triacsin producer, and strongly support that the identified *tri* BGC is related to triacsin biosynthesis since no other homologous BGCs were identified between *S. aureofaciens* and *S. tsukubaensis.* Further sequence analysis showed that the triacsin BGC architecture in *S. tsukubaensis* is nearly identical to that in *S. aureofaciens* except the lack of *tri25,* the gene encoding a putative transposase. By comparing the sequences around the two clusters, the boundary of the *tri* cluster was putatively identified (Figure 1).

Bioinformatic analysis of *tri1-32* (Table S1) revealed that in addition to ORFs that encode a CreE, CreD, and transposase homolog, the cluster encodes eight discrete PKS related proteins: a ketoreductase (KR), three ketosynthases (KS), two dehydratases (DH), an acyl carrier protein (ACP) and a phosphopantetheinyl transferase, which could be involved in the alkyl chain biosynthesis. Seven ORFs were identified to encode oxidoreductases, which may be responsible for regioselective unsaturation of alkyl chains observed in various triacsin congeners as well as the *N*-hydroxytriazene moiety formation. Additional biosynthetic enzymes encoded by the cluster include four ATP-utilizing enzymes, two decarboxylases, two transferase/hydrolase homologs, and three hypothetical proteins. Two putative transcriptional regulators and one transporter encoding genes were also identified in the cluster (Figure 1, Table S1). It is notable that *ŧri26-31* are homologous to the gene cassette *spb38-43* which was essential for the biosynthesis of the glyoxylate hydrazine unit in s56-p1,^[14–15]^ suggesting that at least one N-N bond in triacsins is formed via Tri28, the homolog of Spb40 consisting of a cupin and a methionyl-tRNA synthetase-like protein. In addition, hydrazinoacetic acid (HAA) could be an intermediate for triacsin biosynthesis which is proposed to be generated through the activities of Tri26-28 from lysine and glycine (Figure S8).^[14–15]^ The third nitrogen in the *N*-hydroxytriazene moiety could be derived from nitrous acid due to the presence of Tri16 and 21, the CreD and CreE homologs that presumably generate nitrous acid from aspartate (Figure S8), however enzymes to promote this N-N linkage remain unclear.

In order to probe the necessity of the encoded enzymes in triacsin biosynthesis, we disrupted individual genes for the majority of the ORFs through double crossover in *S. aureofaciens* and/or *S. tsukubaensis* (Figure S3). Triacsin production was completely abolished in *Δtri5-14, 16-19, 21-22, 26-29, 31-32*, demonstrating that these genes Figure 2. uv traces (300 nm) demonstrating the effect of selected were required for triacsin biosynthesis (Figures 2 and S9-10). These results are consistent with the proposed enzymes, including Tri16, 21, 26, 27, 28, which are involved in the *N*-hydroxytriazene moiety formation through two distinct mechanisms for N-N bond formation (Figure S8). Surprisingly, the disruption of *tri23* and *tri24,* encoding two putative saccharopine dehydrogenases, had no effect on any triacsin production in *S. tsukubaensis,* indicating that they are not essential for triacsin biosynthesis (Figure S10). None of the oxidoreductase-encoding gene disruptions led to differential abolishment of triacsin congener production, suggesting that they are not involved in the alkyl chain modification. Additionally, the disruption of *tri3* that encodes one of the four ATP-utilizing enzymes in the cluster decreased triacsin titers by about half in *S. aureofaciens* (Figure 2), suggesting that this gene is also non-essential. The possible role of two regulator-encoding genes, *tri1* and 4, was also examined through gene disruption experiments in *S. aureofaciens. Δtri1* resulted in a small titer increase while *ktrí4* abolished triacsin production (Figure 2), suggesting that the encoded proteins are a negative and a positive transcriptional regulator, respectively.

**Figure 2.**
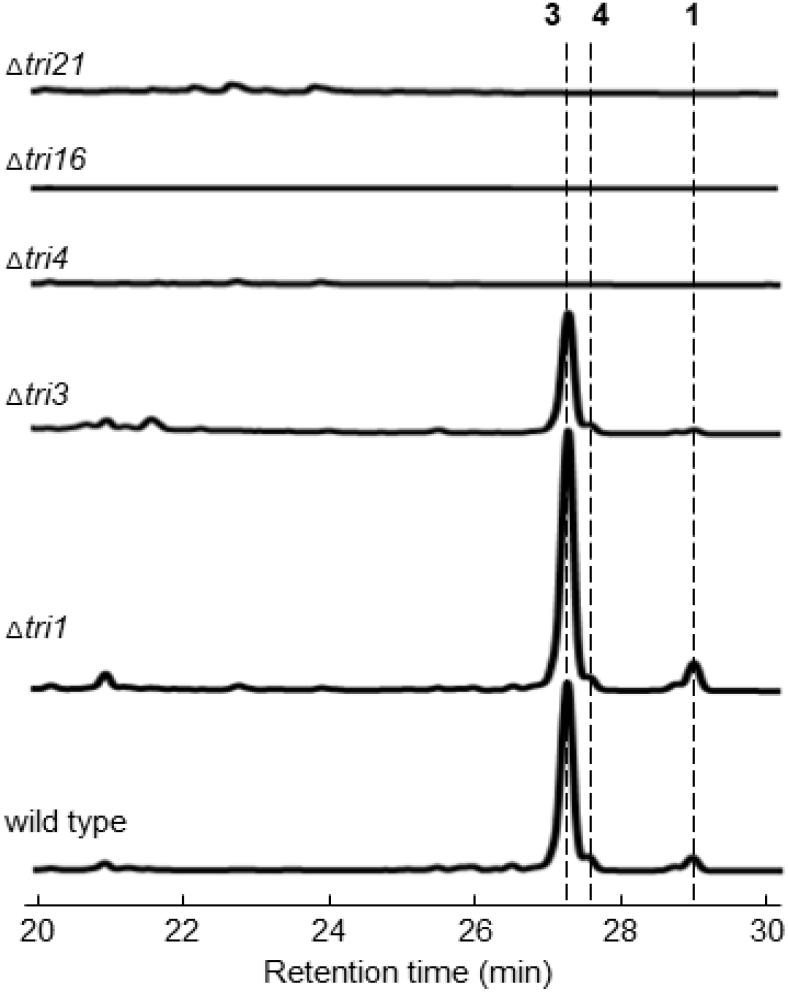
UV traces (300 nm) demonstrating the effect of selected single gene disruptions on triacsin production in *S. aureofaciens*.

To further support the proposed chemical logic behind the biosynthesis of the *N*-hydroxytriazene moiety (Figure S8), we performed labeled precursor feeding studies to probe the utilization of aspartate, nitrite, and glycine in triacsin biosynthesis. Aspartate was proposed to be the substrate for Tri16 and 21 to form nitrous acid and thus one of the nitrogen atoms in triacsins could be derived from the amino group of aspartate. The feeding of ^15^N-aspartate resulted in an enrichment of singly labeled 3 (Figure S11). Additionally, the feeding of ^15^N-nitrite also resulted in an enrichment of singly labeled 3 (Figure S11), consistent with the proposed intermediacy of nitrous acid. Glycine, on the other hand, was proposed to be the substrate for Tri28 to form a hydrazine-containing lysine-glycine adduct, which then undergoes an oxidative cleavage to yield HAA and 2-aminoadipate 6-semialdehyde catalyzed by Tri27, an FAD-dependent D-amino acid oxidase homolog (Figure S8). The feeding of ^15^N-glycine resulted in a significant enrichment of singly labeled 3 (Figure 3A), consistent with the proposed function of Tri28. We next fed 2-^13^C-glycine to further investigate the fate of the α-carbon of glycine in triacsin biosynthesis, considering that all triacsins contain an 11-carbon alkyl chain but PKSs typically generate even-number carbon chains from malonyl-CoAs. The feeding of 2-^13^C-glycine resulted in a weak enrichment of singly labeled 3, far less than the feeding of ^15^N-glycine (Figure 3A). In addition, the enrichment of singly labeled 3 obtained from the feeding of 2-^13^C,^15^N-glycine and ^15^N-glycine was comparable (Figure 3A), making the incorporation of the α-carbon of glycine in triacsin inconclusive. We then scaled up the triacsin-producing cultures supplied with 2-^13^C,^15^N-glycine followed by purification and NMR analysis of 3. Comparison of the ^13^C NMR spectra of 3 produced upon the feeding of 2-^13^C,^15^N-glycine and unlabeled glycine showed that the C1 position was enriched (Figure 3B and S12-13), indicating that this carbon could be derived from the α-carbon of glycine. In addition, splitting of the C1 signal not witnessed in unenriched 3 (Figure 3B) indicates that the N1’ was also enriched, consistent with the hypothesis that the C-N bond of glycine could be incorporated into triacsins as an intact unit, likely via the HAA intermediate (Figure S8). The apparent disproportional ratio of carbon and nitrogen incorporation upon 2-^13^C,^15^N-glycine feeding suggests that glycine molecules could further donate their nitrogen atoms to other positions on triacsins.

**Figure 3:**
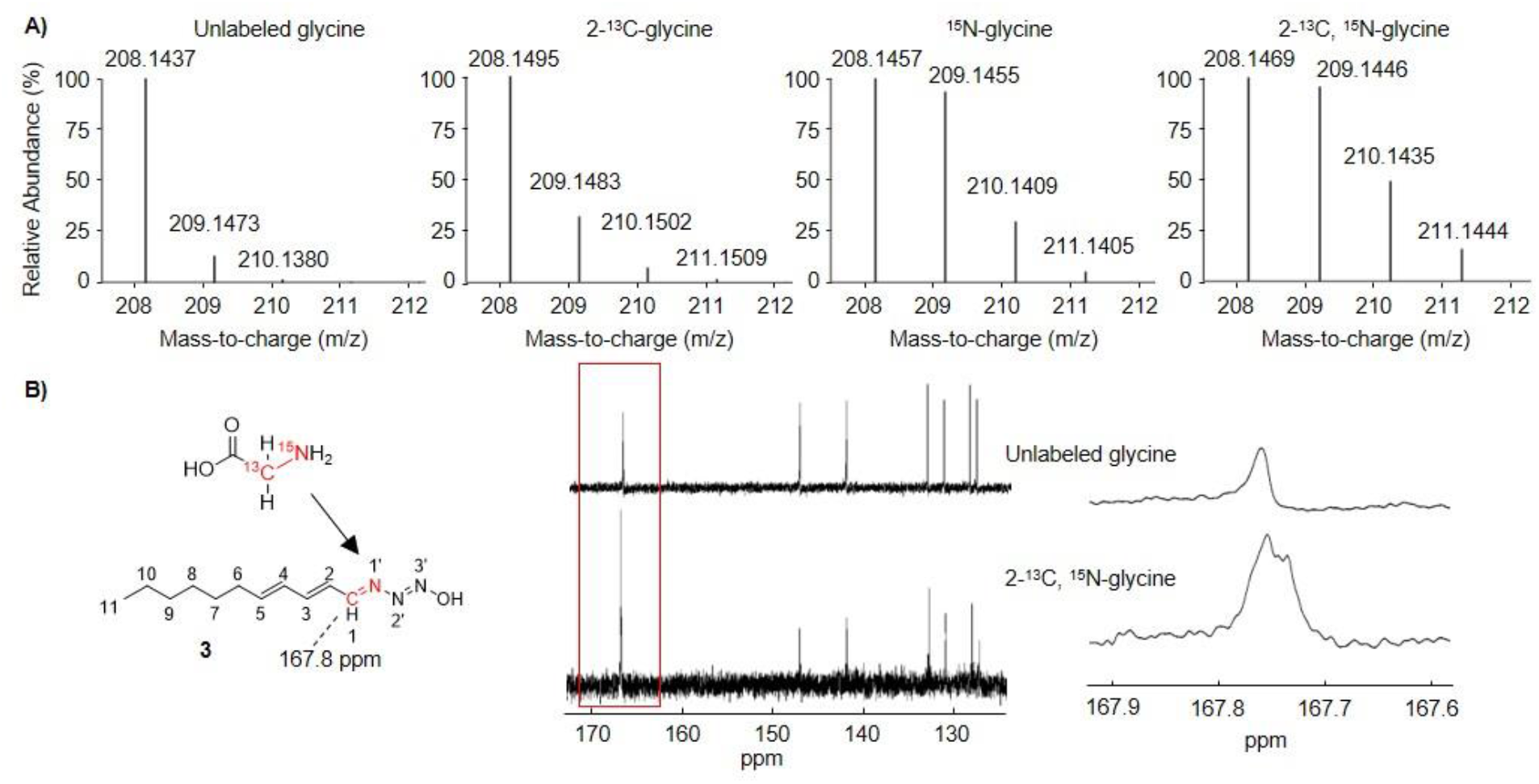
Precursor feeding of isotopically labeled glycine in production of **3**. A) HRMS analysis of 3 from cultures provided with varying isotopically labeled glycine substrates. B) ^13^C NMR spectra of **3** isolated from cultures of *S. aureofaciens* supplied with either 10 mM unlabeled glycine or 2-^13^C, ^15^N-glycine. Close inspection on the enriched C1 signal shows peak splitting, indicative of an adjacent ^15^N, which is not observed in 3 purified from cultures fed unlabeled glycine.

In summary, we have identified the gene cluster for triacsins that sets the stage for further deciphering of the enzymatic machinery for biosynthesis of this class of ACS inhibitors with a unique *N*-hydroxytriazene moiety. Our bioinformatics, mutagenesis, and labeled precursor feeding experiments allow a first proposal of the chemical logic for *N*-hydroxytriazene biosynthesis via an intramolecular reaction of electrophilic and nucleophilic nitrogen as well as a nitrous acid dependent N-N bond formation (Figure S8). The identification of the triacsin gene cluster further led to the discovery of an additional triacsin natural producer. Future biochemical study will provide more insight in the timing and mechanisms of the biosynthesis of this intriguing class of natural products and expand our understanding of enzymes that catalyze the unique *N*-hydroxytriazene moiety formation.

## Experimental Section

Full experimental details are available in the Supporting Information.

## Supporting information

## Acknowledgements

We thank Paul Jensen (the Scripps Institution of Oceanography) for providing *Salinispora arenicola* CNS-205, Jeffrey Skerker (UC Berkeley) for helping with genome assembly, and Jeffrey Pelton for assistance with NMR spectroscopic analysis. This research was financially supported by the National Institutes of Health (DP2AT009148), Alfred P. Sloan Foundation, and the Chan Zuckerberg Biohub investigator program.

