## Supplementary material for "Identifying the Biosynthetic Gene Cluster for Triacsins with an *N*-hydroxytriazene Moiety"

#### Table of Contents

**Experimental Section**

**Supplementary Tables**

**Table S1.** Annotations of the *tri* genes based on sequence homology.
**Table S2.** Plasmids used in this study.
**Table S3.** Primers used in this study.

**Supplementary Figures**

**Figure S1.** Recently examined mechanisms for N-N bond formation.
**Figure S2.** Biosynthetic gene clusters in *S. aureofaciens* encoding CreE and CreD homologs.
**Figure S3.** Gene disruptions in *S. aureofaciens* and *S. tsukubaensis*.
**Figure S4.** HPLC-UV analysis of multigene disruptions in clusters 1 and 2.
**Figure S5.** Gene clusters homologous to the *tri* biosynthetic gene cluster.
**Figure S6.** Production of triacsins by *S. tsukubaensis.*
**Figure S7.** Production of triacsin derivatives by *S. tsukubaensis*.
**Figure S8.** Proposed biosynthetic pathway for triacsin C.
**Figure S9.** HPLC-UV analysis of all gene disruptions in *S. aureofaciens.*
**Figure S10.** HPLC-UV analysis of all gene disruptions in *S. tsukubaensis*.
**Figure S11.** LC-MS isotopic peak analysis of **3** after $^{15}$N-nitrite and $^{15}$N-aspartate precursor feeding studies.
**Figure S12.** NMR characterization of **3** purified from cultures fed unlabeled glycine.
**Figure S13.** NMR characterization of **3** purified from cultures fed 2-$^{13}$C, $^{15}$N-glycine.

### Experimental Section

**Bacterial Strains and Growth Conditions.** *Streptomyces aureofaciens* ATCC 31442 was cultured on ISP4 agar plates to allow sporulation. Individual spore colonies would be inoculated into a 2 mL seed culture of R5 liquid media and grown at 30°C and 240 rpm for 48 hours. The seed culture would be inoculated into 30 mL R5 liquid media and grown for 5 days at 30°C and 240 rpm before cell harvesting. Culturing was performed in 125 mL Erlenmeyer vessels with a wire coil placed at the bottom to encourage aeration. *Streptomyces tsukubaensis* NRRL 18488 was cultured using the same methodology with the exception that, in place of R5 media, we used ISP2 liquid media buffered with 5 g/L TES free acid and titrated to a final pH of 7.4. For all PCR targeting with the λ RED recombination system we used *E. coli* BW25113 with pIJ790. *E. coli* Transformax EPI300 (Epicentre) were used for the generation of a fosmid library and the maintenance of large fosmids. *E. coli* One Shot TOP10 (ThermoFisher Scientific) was used for ligation independent cloning using the Zero Blunt TOPO Cloning Kit (ThermoFisher Scientific). For all other routine cloning procedures, we used *E. coli* XL1-Blue. Conjugations to transfer genetic material to streptomycetes was performed with *E. coli* WM6026 which requires an exogeneous supply of diaminopimelic acid (DAP). All *E. coli* were cultured in LB at 37°C unless otherwise specified. Growth media was supplied with antibiotics as required at the following concentrations: kanamycin (50 µg/mL), apramycin (50 µg/mL), ampicillin (100 µg/mL), and chloramphenicol (25 µg/mL).

**Construction of Fosmids for Gene Disruption in *S. aureofaciens* ATCC 31442.** Genomic DNA from *S. aureofaciens* was purified using the Quick-DNA Fungal/Bacterial Kit (Zymo Research) and sent to the University of California, Davis Genome Center for Illumina and PacBio sequencing. A fosmid library was generated through the CopyControl HTP Fosmid Library Production Kit (Epicentre) with the pCC2FOS vector. The library was screened using primers designed to PCR amplify fosmids containing targeted genes. Junctions between the pCC2FOS vector and the insertion were confirmed by DNA sequencing (UC Berkeley DNA Sequencing Facility). The fosmids were purified using the ZR BAC DNA Miniprep Kit (Zymo Research) and transformed into *E. coli* BW25113 for PCR targeting. Gene disruption plasmids were then generated by standard procedure using pIJ773 as a template for disruption cassettes conferring apramycin resistance. Resulting gene disruption fosmids were propagated in *E. coli* Transformax EPI300 and then confirmed by restriction digest and DNA sequencing (UC Berkeley DNA Sequencing Facility)

**Construction of Plasmids for Gene Disruption in *S. tsukubaenesis* NRRL 18488.** Genomic DNA from *S. tsukubaensis* was purified using the Quick-DNA Fungal/Bacterial Kit (Zymo Research). Regions spanning 6 – 10 kb were PCR amplified with PrimeSTAR GXL DNA Polymerase (Takara) and ligated into the pCR-Blunt II-TOPO vector (ThermoFisher Scientific). The gene disruption plasmids were made by an identical procedure to generation of gene disruption fosmids with the exception that *E. coli* XL1-Blue was used in place of *E. coli* Transformax EPI300 for plasmid propagation.

**Gene Disruptions in *S. aureofaciens* and *S. tsukubaensis*.** PCR-targeting gene disruption fosmids or plasmids were transformed into *E. coli* WM6026. *E. coli* WM6026 was grown in 10 mL LB at 37°C supplied with 0.5 mM DAP. Once the $OD_{600}$ had reached 0.6, the culture was pelleted by centrifugation (4,000 x g, 5 min) at 4°C. After decanting the media, the pellet was resuspended in 10 mL cold LB for two additional washes to remove residual antibiotics and DAP. During this preparation of *E. coli* WM6026, spores from the recipient streptomycete were harvested from an ISP4 agar plate in a 10% glycerol solution. The spore pellet was pelleted by centrifugation (10,000 x g, 30 sec), decanted, and then resuspended in 500 µL LB. The spore solution was heat shocked at 50°C for 10 min and then pelleted by centrifugation as before. After decanting the media, the spore pellet was resuspended in 300 µL LB. This spore solution was then used to resuspend the *E. coli* WM6026 pellet with the desired gene disruption construct. The mixed spore/*E. coli* solution was plated on ISP4 agar plates supplemented with an additional 10 mM calcium chloride and 10 mM magnesium chloride. After a period of 18 hours, the plates were overlaid in an aqueous solution containing 3.5 mg of apramycin. After 4 – 7 days, exconjugants were replica plated onto ISP4 plates containing either apramycin or kanamycin. Double crossover mutants displaying kanamycin sensitivity and apramycin resistance were cultured and both the presence and position of the apramycin gene disruption cassette were confirmed by PCR. If only single crossover mutants resistant to both selective agents were obtained, then the mutants were sub-cultivated on ISP4 without antibiotics and this process was repeated until double crossover mutants were isolated.

**LC-MS Analysis of Triacsins.** Streptomycete cultures were pelleted by centrifugation (4,000 x g for 15 min), and the spent media was extracted 1:1 by volume with ethyl acetate. The ethyl acetate was removed by rotary evaporation, and the dry extract was dissolved in 300 µL of methanol for liquid chromatography-mass spectrometry (LC-MS) analysis by 10 µL injection onto an Agilent Technologies 6120 Quadrupole LC-MS instrument with an Agilent Eclipse Plus C18 column (4.6 x 100 mm). For high resolution MS (HRMS) analysis, the extract in methanol could be diluted by a factor of 5 for LC-HRMS and HRMS/MS analysis by 10 µL injection onto an Agilent Technologies 6510 Q-TOF LC-MS instrument with the same column. Linear gradients of 5-95% acetonitrile (vol/vol) in water with 0.1% formic acid (vol/vol) at 0.5 ml/min were used.

**Production and Purification of Triacsins.** After the 5-day culture period for *S. aureofaciens*, the cultures were pelleted by centrifugation (4,000 x g for 15 min), and the spent media was extracted 1:1 by volume with ethyl acetate. The ethyl acetate was removed by rotary evaporation, and the dry extract was dissolved in a 1:1 mixture (vol/vol) of dichloromethane and methanol. The dissolved extract was loaded onto a size exclusion column packed with Sephadex LH-20 (Sigma-Aldrich) and manually fractionated in a 1:1 dichloromethane and methanol running solvent. The fractions were screened by LC-MS with a diode array detector (DAD). Fractions containing triacsins were subjected to high performance liquid chromatography (HPLC) using an Agilent 1260 HPLC and an Atlantis T3 OBD Prep column (100 Å pore size, 5 µm particle size, 10 x 150 mm). Linear gradients of 5-95% acetonitrile (vol/vol) in water with 0.1% formic acid (vol/vol) at 0.5 ml/min were used. Purified products were dried and analyzed by LC-MS and NMR. All NMR spectra were recorded on a Bruker AVANCE at 900 MHz ($^{1}$H NMR) and 226 MHz ($^{13}$C NMR).

**Labeled Precursor Feeding Experiments.** *S. aureofaciens* was cultured as described above and 24 hours after the 30 mL culture inoculation, the culture was supplied with glycine or aspartate to a final concentration of 10 mM. Substrates tested were unlabeled glycine, 2-$^{13}$C-glycine, $^{15}$N-glycine, 2-$^{13}$C,$^{15}$N-glycine, and $^{15}$N-aspartate. The same methodology was used for supplying liquid cultures of *S. aureofaciens* with either sodium nitrite or sodium $^{15}$N-nitrite, but a final concentration of 1 mM was used. Compound extraction and LC-HRMS analysis were performed as described above.

### Supplementary Tables

**Table S1.** Annotations of the *tri* genes based on sequence homology.

| Gene | Size (aa) | Annotation | Protein Homolog Origin* | Accession Number | Identity (%) | Similarity (%) |
|---|---|---|---|---|---|---|
| *tri1* | 225 | DNA-binding response regulator | *Streptomyces clavuligerus* | WP_003956729 | 98 | 99 |
| *tri2* | 498 | MFS transporter | *Streptomyces clavuligerus* | WP_003956728.1 | 77 | 88 |
| *tri3* | 522 | long-chain fatty acid--CoA ligase | *Streptomyces yerevanensis* | WP_033320718.1 | 76 | 85 |
| *tri4* | 462 | transciptional regulator | *Streptomyces ipomoeae* | WP_009323372.1 | 69 | 81 |
| *tri5* | 234 | ketoreductase | *Streptomyces ipomoeae* | WP_048821009.1 | 83 | 90 |
| *tri6* | 382 | ketosynthase | *Streptomyces ipomoeae* | WP_048821010.1 | 83 | 87 |
| *tri7* | 186 | ketosynthase | *Streptomyces ipomoeae* | WP_009323339.1 | 75 | 82 |
| *tri8* | 225 | hypothetical protein | *Streptomyces ipomoeae* | WP_009323369.1 | 75 | 90 |
| *tri9* | 247 | UbiX family flavin prenyltransferase | *Streptomyces ipomoeae* | WP_009323418.1 | 84 | 93 |
| *tri10* | 490 | UbiD family decarboxylase | *Streptomyces ipomoeae* | WP_009323364.1 | 79 | 87 |
| *tri11* | 154 | MaoC family dehydratase | *Streptomyces ipomoeae* | WP_009323341.1 | 81 | 87 |
| *tri12* | 143 | MaoC family dehydratase | *Streptomyces ipomoeae* | WP_009323416.1 | 79 | 82 |
| *tri13* | 310 | ketosynthase | *Salinispora pacifica* | WP_018723633.1 | 73 | 84 |
| *tri14* | 409 | peptidase C45 | *Streptomyces ipomoeae* | WP_009318533.1 | 73 | 82 |
| *tri15* | 249 | ppant transferase | *Streptomyces ipomoeae* | WP_009318532.1 | 58 | 66 |
| *tri16* | 565 | nitrosuccinate lyase | *Salinispora pacifica* | WP_025617508.1 | 70 | 76 |
| *tri17* | 555 | long-chain fatty acid--CoA ligase | *Streptomyces ipomoeae* | WP_048820752.1 | 80 | 87 |
| *tri18* | 498 | oxidoreductase | *Streptomyces ipomoeae* | WP_048820362.1 | 64 | 81 |
| *tri19* | 405 | oxidoreductase | *Streptomyces ipomoeae* | WP_009312596.1 | 65 | 76 |
| *tri20* | 81 | acyl carrier protein | *Streptomyces ipomoeae* | WP_009312594.1 | 69 | 82 |
| *tri21* | 645 | flavin-dependent monooxygenase | *Salinispora pacifica* | WP_018723641.1 | 67 | 75 |
| *tri22* | 399 | acyl-CoA dehydrogenase | *Streptomyces ipomoeae* | WP_078613912.1 | 79 | 88 |
| *tri23* | 386 | saccharopine dehydrogenase | *Streptomyces ochraceiscleroticus* | WP_031057382.1 | 66 | 70 |
| *tri24* | 429 | saccharopine dehydrogenase | *Streptomyces ipomoeae* | WP_009311547.1 | 68 | 74 |

| Gene | Size (aa) | Annotation | Protein Homolog Origin | Accession Number | Identity (%) | Similarity (%) |
|---|---|---|---|---|---|---|
| *tri25* | 390 | transposase | *Streptomyces orinoci* | WP_109284379.1 | 87 | 92 |
| *tri26* | 439 | lysine monooxygenase | *Streptomyces ipomoeae* | WP_009311506.1 | 81 | 89 |
| *tri27* | 358 | oxidoreductase | *Streptomyces ipomoeae* | WP_009311508.1 | 70 | 78 |
| *tri28* | 678 | cupin-domain containing tRNA synthetase | *Streptomyces ipomoeae* | WP_009311527.1 | 78 | 84 |
| *tri29* | 513 | AMP-dependent synthetase | *Streptomyces ipomoeae* | WP_009311533.1 | 74 | 83 |
| *tri30* | 90 | acyl carrier protein | *Streptomyces ipomoeae* | WP_009311519.1 | 84 | 87 |
| *tri31* | 195 | N-acetyltransferase | *Streptomyces ipomoeae* | WP_009311512.1 | 85 | 91 |
| *tri32* | 168 | DUF4033 domain-containing protein | *Nocardia sp. NRRL S-836* | WP_053732975.1 | 35 | 48 |

*All hits are the highest-ranking excluding those from *S. tsukubaensis*.

**Table S2.** Plasmids used in this study.

| **Plasmid** | **Derived from** | **Function** |
|---|---|---|
| pCC2FOS-cluster1 | pCC2FOS | 9 kb gene disruption in cluster 1 |
| pCC2FOS-cluster2 | pCC2FOS | 9 kb gene disruption in cluster 2, *tri* biosynthetic gene cluster |
| pCC2FOS-*tri1* | pCC2FOS | Gene disruption of *tri1* in *Streptomyces aureofaciens* |
| pCC2FOS-*tri3* | pCC2FOS | Gene disruption of *tri3* in *Streptomyces aureofaciens* |
| pCC2FOS-*tri4* | pCC2FOS | Gene disruption of *tri4* in *Streptomyces aureofaciens* |
| pCC2FOS-*tri6* | pCC2FOS | Gene disruption of *tri6* in *Streptomyces aureofaciens* |
| pCC2FOS-*tri9*-*tri10* | pCC2FOS | Gene disruption of *tri9-10* in *Streptomyces aureofaciens* |
| pCC2FOS-*tri14* | pCC2FOS | Gene disruption of *tri14* in *Streptomyces aureofaciens* |
| pCC2FOS-*tri16* | pCC2FOS | Gene disruption of *tri16* in *Streptomyces aureofaciens* |
| pCC2FOS-*tri18* | pCC2FOS | Gene disruption of *tri18* in *Streptomyces aureofaciens* |
| pCC2FOS-*tri21* | pCC2FOS | Gene disruption of *tri21* in *Streptomyces aureofaciens* |
| pCR-Blunt-*tri5* | pCR-Blunt II-TOPO | Gene disruption of *tri5* in *Streptomyces tsukubaensis* |
| pCR-Blunt-*tri7* | pCR-Blunt II-TOPO | Gene disruption of *tri7* in *Streptomyces tsukubaensis* |
| pCR-Blunt-*tri8* | pCR-Blunt II-TOPO | Gene disruption of *tri8* in *Streptomyces tsukubaensis* |
| pCR-Blunt-*tri11* | pCR-Blunt II-TOPO | Gene disruption of *tri11* in *Streptomyces tsukubaensis* |
| pCR-Blunt-*tri12* | pCR-Blunt II-TOPO | Gene disruption of *tri12* in *Streptomyces tsukubaensis* |
| pCR-Blunt-*tri13* | pCR-Blunt II-TOPO | Gene disruption of *tri13* in *Streptomyces tsukubaensis* |
| pCR-Blunt-*tri17* | pCR-Blunt II-TOPO | Gene disruption of *tri17* in *Streptomyces tsukubaensis* |
| pCR-Blunt-*tri19* | pCR-Blunt II-TOPO | Gene disruption of *tri19* in *Streptomyces tsukubaensis* |
| pCR-Blunt-*tri22* | pCR-Blunt II-TOPO | Gene disruption of *tri22* in *Streptomyces tsukubaensis* |
| pCR-Blunt-*tri23* | pCR-Blunt II-TOPO | Gene disruption of *tri23* in *Streptomyces tsukubaensis* |
| pCR-Blunt-*tri24* | pCR-Blunt II-TOPO | Gene disruption of *tri24* in *Streptomyces tsukubaensis* |
| pCR-Blunt-*tri26* | pCR-Blunt II-TOPO | Gene disruption of *tri26* in *Streptomyces tsukubaensis* |
| pCR-Blunt-*tri27* | pCR-Blunt II-TOPO | Gene disruption of *tri27* in *Streptomyces tsukubaensis* |
| pCR-Blunt-*tri28* | pCR-Blunt II-TOPO | Gene disruption of *tri28* in *Streptomyces tsukubaensis* |
| pCR-Blunt-*tri29* | pCR-Blunt II-TOPO | Gene disruption of *tri29* in *Streptomyces tsukubaensis* |
| pCR-Blunt-*tri31* | pCR-Blunt II-TOPO | Gene disruption of *tri31* in *Streptomyces tsukubaensis* |
| pCR-Blunt-*tri32* | pCR-Blunt II-TOPO | Gene disruption of *tri32* in *Streptomyces tsukubaensis* |

**Table S3.** Primers used in this study.

| Primer | Sequence (5' → 3') | Description |
| --- | --- | --- |
| cluster1-F | tcgacgacgaccacccctctatctccagtacacctATGATTCCGGGGATCCGTCGACC | 9 kb multigene disruption in cluster 1 |
| cluster1-R | cccggccgatcctcccatactctccacttgccgataTCATGTAGGCTGGAGCTGCTTC | |
| cluster2-F | atcacatcgaagaacgccaggccgaggccgcgcaggATGATTCCGGGGATCCGTCGACC | 9 kb multigene disruption in cluster 2 |
| cluster2-R | caacgggaacctggacgaccagacctgtgtgctctcTCATGTAGGCTGGAGCTGCTTC | |
| *tri1*-F | atgacgcgtgtactgctcgcagaggacgacgcatccATGATTCCGGGGATCCGTCGACC | Gene disruption of *tri1* in *Streptomyces aureofaciens* |
| *tri1*-R | tacatcgcgacggtccgcggcgtcggcttccgcttcTCATGTAGGCTGGAGCTGCTTC | |
| *tri3*-F | gaggtgcggggcgtggcgaaagggctgatcgcctcgATGATTCCGGGGATCCGTCGACC | Gene disruption of *tri3* in *Streptomyces aureofaciens* |
| *tri3*-R | gcaccgtcgtgtccggacgccgtggggcggggccggTCATGTAGGCTGGAGCTGCTTC | |
| *tri4*-F | gccacaggaacctttcgaccgagaggcaccacgttcATGATTCCGGGGATCCGTCGACC | Gene disruption of *tri4* in *Streptomyces aureofaciens* |
| *tri4*-R | gaagccctggcctccgactcggctcccgccgtgacgTCATGTAGGCTGGAGCTGCTTC | |
| *tri6*-F | ttcaagcccgtgaccaggttcgacaccgaccggtacATGATTCCGGGGATCCGTCGACC | Gene disruption of *tri6* in *Streptomyces aureofaciens* |
| *tri6*-R | gtcgagggccagtaccgagatgaccagctcgaccatTCATGTAGGCTGGAGCTGCTTC | |
| *tri9-10*-F | cgccgaccgaccgaattccgccgaaggagcagcccaATGATTCCGGGGATCCGTCGACC | Gene disruption of *tri9-10* in *Streptomyces aureofaciens* |
| *tri9-10*-R | tcggtgggcagtcagtcgtcccagtgctccagcaccTCATGTAGGCTGGAGCTGCTTC | |
| *tri14*-F | ccgcaagtgcccgtgacgggaaggggtgtgccggacATGATTCCGGGGATCCGTCGACC | Gene disruption of *tri14* in *Streptomyces aureofaciens* |
| *tri14*-R | ggtccgctcccgccggatgttcccgttgtcgggaacTCATGTAGGCTGGAGCTGCTTC | |
| *tri16*-F | gtggtgtcggaccgggcgtggctcgacgggatggtcATGATTCCGGGGATCCGTCGACC | Gene disruption of *tri16* in *Streptomyces aureofaciens* |
| *tri16*-R | ttccgcggcgcgtacggcacgcgtcacgatctgtttTCATGTAGGCTGGAGCTGCTTC | |
| *tri18*-F | ggcctcgaccgtgattcctccggagggagacaacccATGATTCCGGGGATCCGTCGACC | Gene disruption of *tri18* in *Streptomyces aureofaciens* |
| *tri18*-R | tcacagcacctccagcagccgccgggacatccgggcTCATGTAGGCTGGAGCTGCTTC | |
| *tri21*-F | atgtcacagccgcctcggacggtcgcggtcgtgggcATGATTCCGGGGATCCGTCGACC | Gene disruption of *tri21* in *Streptomyces aureofaciens* |
| *tri21*-R | tcaccggccgagcaccgctctggccaccgcgtcggcTCATGTAGGCTGGAGCTGCTTC | |
| *tri5*-F | tgctgttttccacgaccaccgaccgggctggagtttgttcATTCCGGGGATCCGTCGACC | Gene disruption of *tri5* in *Streptomyces tsukubaensis* |
| *tri5*-R | tcccggccgggtattccccgggtttttccggcccctccatgTGTAGGCTGGAGCTGCTTC | |
| *tri7*-F | gcggcgcgaacacggcgctggtggtgagcgcaccatgaccATTCCGGGGATCCGTCGACC | Gene disruption of *tri7* in *Streptomyces tsukubaensis* |
| *tri7*-R | gacgcccagttcgaccagcaccaccaggacgtcccgggcgcTGTAGGCTGGAGCTGCTTC | |
| *tri8*-F | gggcgcggaaggggtggcggaatgagggagcgcgccgtggATTCCGGGGATCCGTCGACC | Gene disruption of *tri8* in *Streptomyces tsukubaensis* |
| *tri8*-R | gcgcatacaggccacgacctcgtcgtccgccgtgacgccgaTGTAGGCTGGAGCTGCTTC | |
| *tri11*-F | gttccgtgcccctccggggccttcgaccgaaggaagcgatATTCCGGGGATCCGTCGACC | Gene disruption of *tri11* in *Streptomyces tsukubaensis* |
| *tri11*-R | gcgctggttgccgtggatcagatacgtggggtcgaagccgaTGTAGGCTGGAGCTGCTTC | |
| *tri12*-F | aaggaggtgaacccgcatgaccacggacacggcaggggcgATTCCGGGGATCCGTCGACC | Gene disruption of *tri12* in *Streptomyces tsukubaensis* |
| *tri12*-R | gaccgcgcgggcttcgctcagcaccttcgtaccgccgcaggTGTAGGCTGGAGCTGCTTC | |
| *tri13*-F | ctgcccgtgcacggcgatcccgaacggagccccaggacggATTCCGGGGATCCGTCGACC | Gene disruption of *tri13* in *Streptomyces tsukubaensis* |
| *tri13*-R | gtcgtccgccgcctctccagtggcgagcagatcggccaggtTGTAGGCTGGAGCTGCTTC | |
| *tri17*-F | gaaggtgctcgccgacgcggcgctcggcgcggggaacgtgATTCCGGGGATCCGTCGACC | Gene disruption of *tri17* in *Streptomyces tsukubaensis* |
| *tri17*-R | caggcgagccccggcccgctgccgcggtcgtacgggacggaTGTAGGCTGGAGCTGCTTC | |
| *tri19*-F | tctcatccgtactgtccggctccgggaacgaggtccactgATTCCGGGGATCCGTCGACC | Gene disruption of *tri19* in *Streptomyces tsukubaensis* |
| *tri19*-R | gtttctcgggtgtcccggttttcccgggtgtcccgggaaacTGTAGGCTGGAGCTGCTTC | |

| Primer | Sequence (5' → 3') | Description |
| --- | --- | --- |
| *tri22*-F | cccccttctacgacgacgggcaccggcgcctggccggcgagATTCCGGGGATCCGTCGACC | Gene disruption of *tri22* in *Streptomyces tsukubaensis* |
| *tri22*-R | tccggacaggggttccggacaggggttccggtgccgggccgTGTAGGCTGGAGCTGCTTC | |
| *tri23*-F | gcccggccgacgcgccccggcccggcgaggagcagcactgATTCCGGGGATCCGTCGACC | Gene disruption of *tri23* in *Streptomyces tsukubaensis* |
| *tri23*-R | gaagccgcggctcgcctcggcgggcagcagcgacggcagatTGTAGGCTGGAGCTGCTTC | |
| *tri24*-F | cacgagcggtgtcgtccactgggtggggaccggtctctccATTCCGGGGATCCGTCGACC | Gene disruption of *tri24* in *Streptomyces tsukubaensis* |
| *tri24*-R | taccacccccagcgccagcggcagcgacacacagcgggccaTGTAGGCTGGAGCTGCTTC | |
| *tri26*-F | gcgcagttcgacgccgtcccgcgctccgagttccgcaactATTCCGGGGATCCGTCGACC | Gene disruption of *tri26* in *Streptomyces tsukubaensis* |
| *tri26*-R | cacgtccgcctcgaacagttcggtggagtccggatggtcgtTGTAGGCTGGAGCTGCTTC | |
| *tri27*-F | cgaacccggcacctcggtgaccgtcctggaccgggagctgATTCCGGGGATCCGTCGACC | Gene disruption of *tri27* in *Streptomyces tsukubaensis* |
| *tri27*-R | gtcgccggtcagcggccggaccagcggtcctccggccgggtTGTAGGCTGGAGCTGCTTC | |
| *tri28*-F | gaaccgggtgtcctcgccctcttcgaacccttcgagacccATTCCGGGGATCCGTCGACC | Gene disruption of *tri28* in *Streptomyces tsukubaensis* |
| *tri28*-R | cctgcccgtcggtcccgtcggtgccgtccgcaccggtgcttTGTAGGCTGGAGCTGCTTC | |
| *tri29*-F | agaccgcgggcttcgaacccggcaagttcacccccggtggATTCCGGGGATCCGTCGACC | Gene disruption of *tri29* in *Streptomyces tsukubaensis* |
| *tri29*-R | cagggcctcctccaccggccggaccacgaccttcttcccgtTGTAGGCTGGAGCTGCTTC | |
| *tri31*-F | atctggcggtacttcgtggccgccgtccacgacgacaacgATTCCGGGGATCCGTCGACC | Gene disruption of *tri31* in *Streptomyces tsukubaensis* |
| *tri31*-R | gagccgttccttcacgcccggccactccgccttcaggatgcTGTAGGCTGGAGCTGCTTC | |
| *tri32*-F | cctcgtgttcgactccctgcttctgaagaagttcaacaggATTCCGGGGATCCGTCGACC | Gene disruption of *tri32* in *Streptomyces tsukubaensis* |
| *tri32*-R | cgccgccgttcacgcctccggagcgcggcgctccggaggcgTGTAGGCTGGAGCTGCTTC | |
| T7 primer | taatacgactcactataggg | Sequencing pCC2FOS junctions |
| pCC2FOS-R | gcttgcatgcctgcaggt | |
| M13-F | gtaaaacgacggccag | Sequencing pCR-Blunt II-TOPO junctions |
| M13-R | caggaaacagctatgac | |

### Supplementary Figures

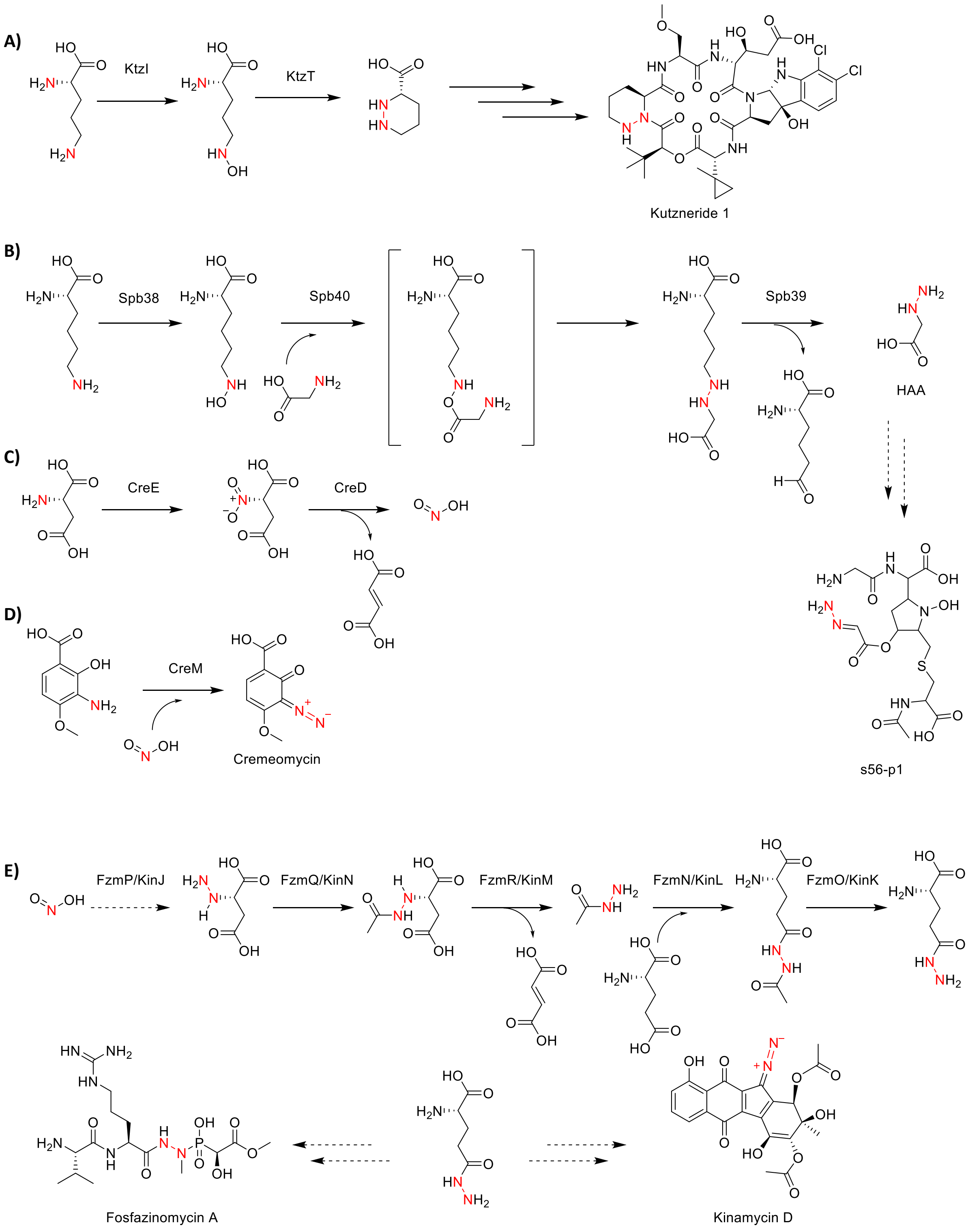


**Figure S1.** Recently examined mechanisms for N-N bond formation. A) KtzI and KtzT catalyze the formation of piperazic acid, a non-proteinogenic amino acid incorporated in several nonribosomal peptides such as the kutznerides. B) Three enzymes encoded in the *spb* BGC have been shown to form HAA from an N-N ligation product of lysine and glycine. C) Nitrous acid produced from aspartate through the action of CreE and CreD. Nitrous acid production by homologs of these two enzymes has been implicated in several BGCs of known N-N bond-containing natural products. D) In the cremeomycin biosynthetic pathway CreM catalyzes the diazotization of the penultimate cremeomycin precursor by utilizing ATP to activate nitrous acid. E) In both the fosfazinomycin and kinamycin pathways, a gene cassette involved in the assembly and transfer of hydrazine using glutamate as a carrier has been characterized. However the dedicated N-N forming enzymes using nitrous acid have not been identified in these cases.

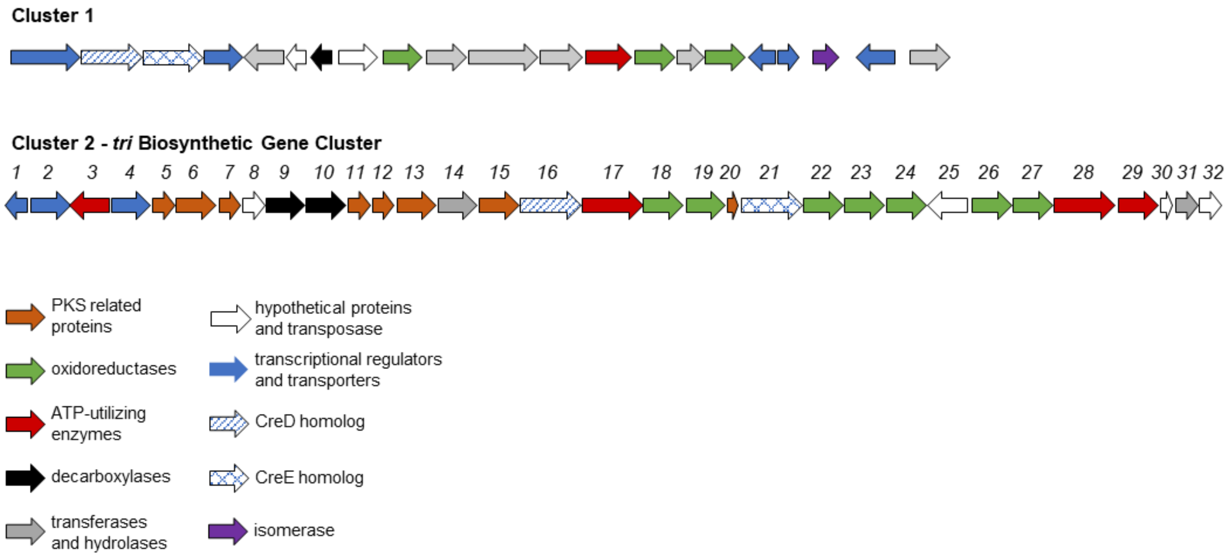


**Figure S2.** Biosynthetic gene clusters in *S. aureofaciens* encoding CreE and CreD homologs.

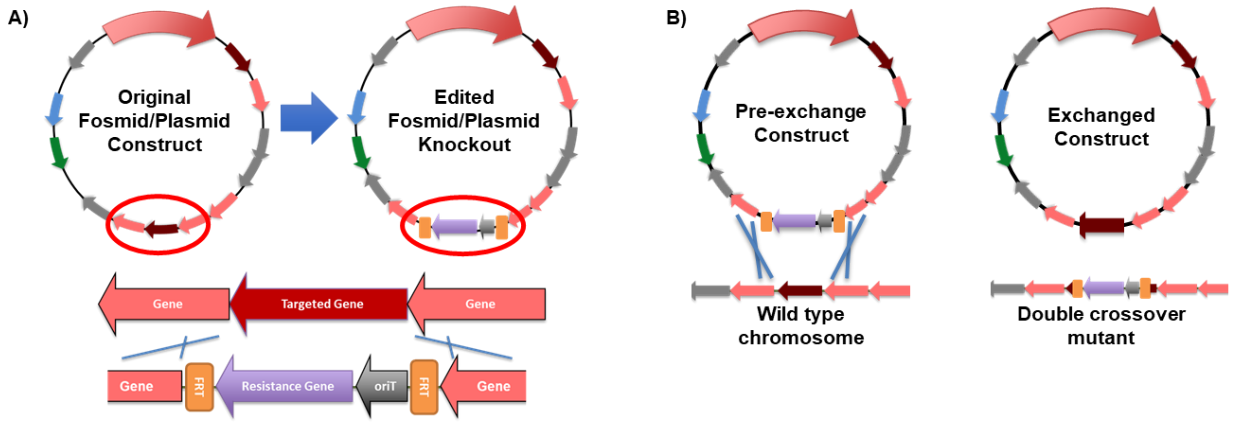


C)

| Expected mutant size | Gene Disrupted |
|---|---|
| 1723 | Δ*tri1* |
| 1440 | Δ*tri3* |
| 1008 | Δ*tri6* |
| 1829 | Δ*tri9-10* |
| 1577 | Δ*tri16* |
| 1112 | Δ*tri21* |

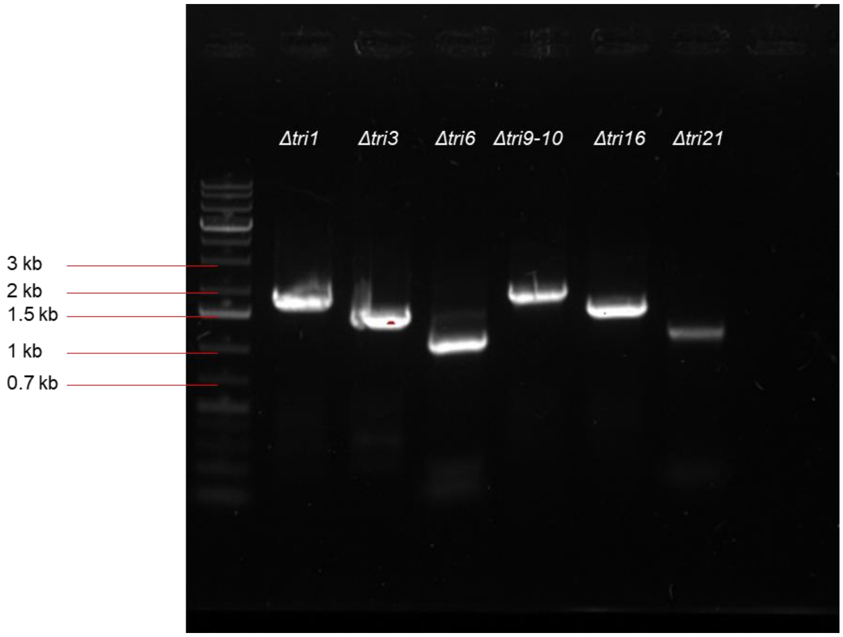


**Figure S3.** Gene disruptions in *S. aureofaciens* and *S. tsukubaensis*. All gene disruptions were performed through homologous recombination. A) Fosmid (*S. aureofaciens*) and plasmid (*S. tsukubaensis*) disruption constructs were created through the λ RED recombination system in *E. coli* BW25113 bearing pIJ790 to replace the targeted ORF with a selective marker. B) Constructs were transferred to the streptomycete recipient by conjugation using *E. coli* WM6026 and double crossover exconjugants were screened as described in the experimental section. C) A sample gel confirming the location of the disruption cassettes in mutants. Genomic DNA was extracted from the mutants and confirmed via PCR using one primer internal to the disruption cassette, and one external in the flanking genes. The sizes of the amplified PCR products were consistent with the expected sizes for the mutants.

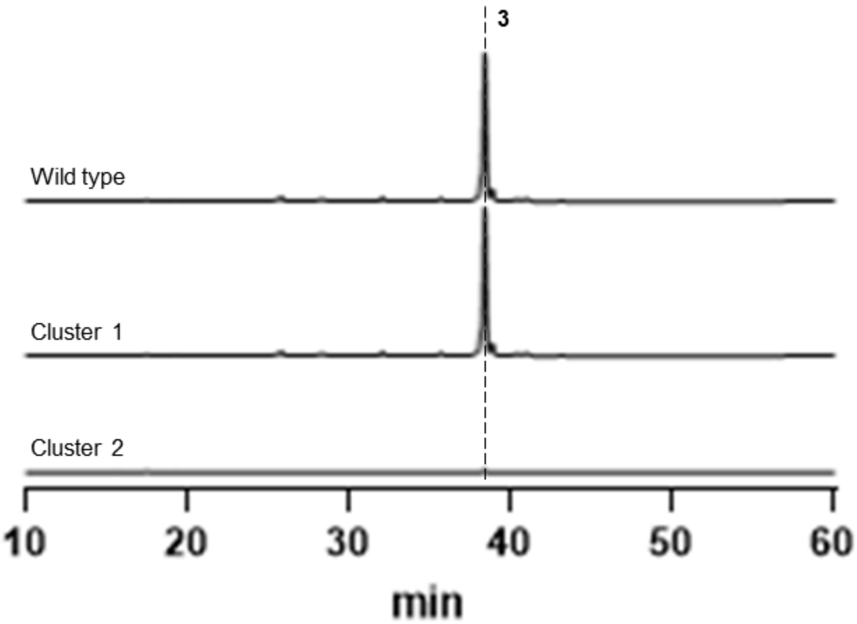


**Figure S4.** HPLC-UV analysis (300 nm) of multigene disruptions in clusters 1 and 2.

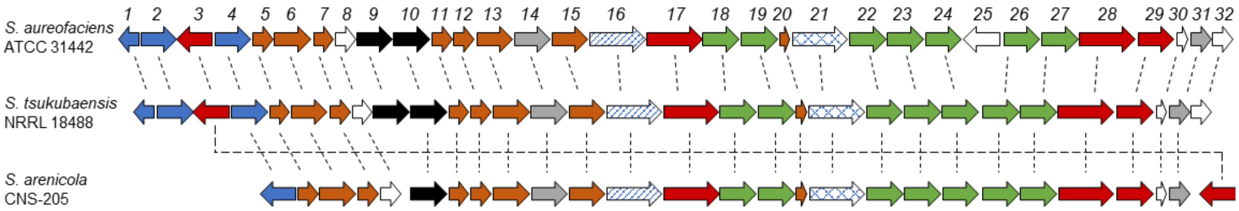


**Figure S5.** Gene clusters homologous to the *tri* biosynthetic gene cluster.

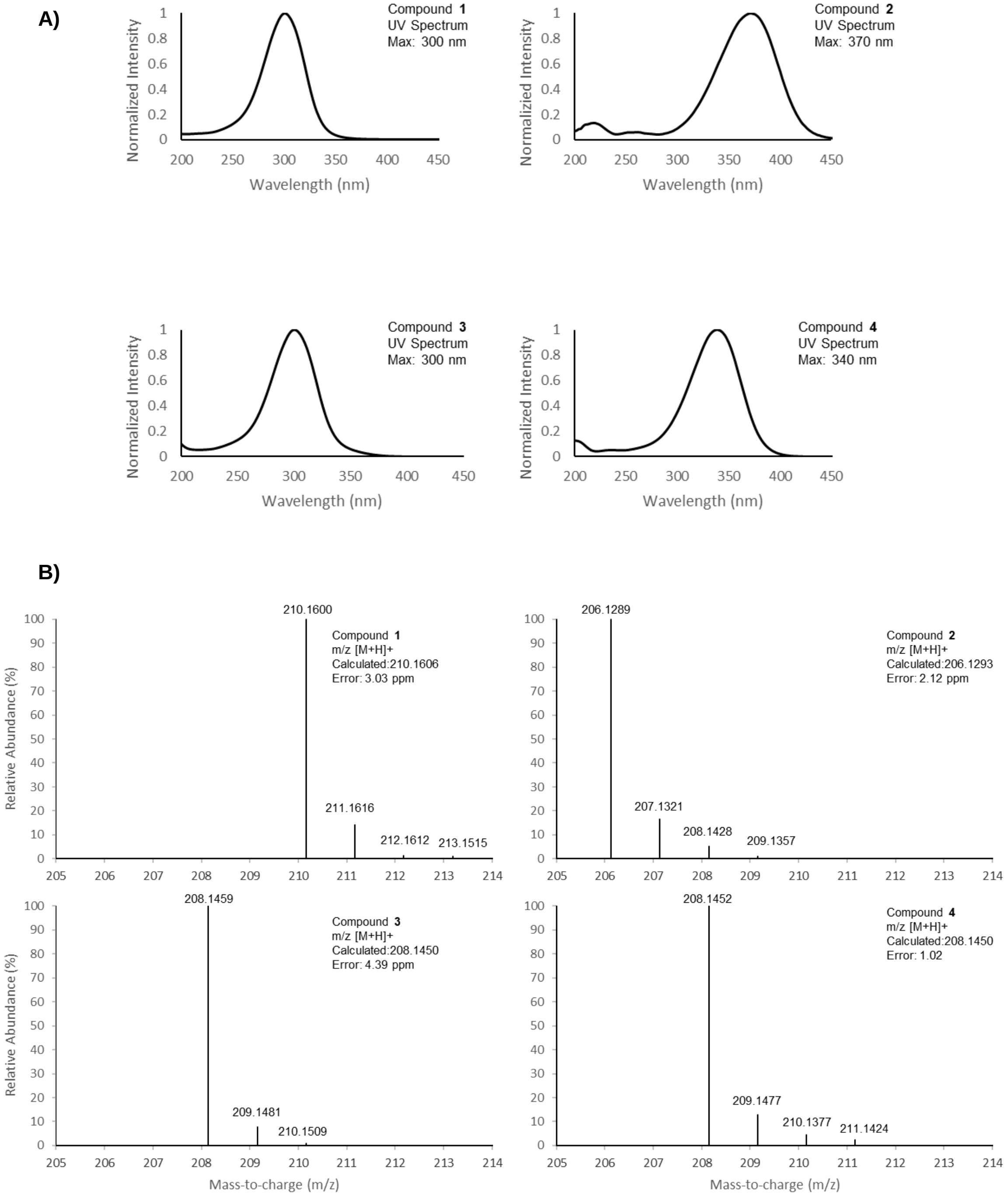


**Figure S6.** Production of triacsins by *S. tsukubaensis*. A) UV spectra for **1-4**. B) HRMS for **1-4**.

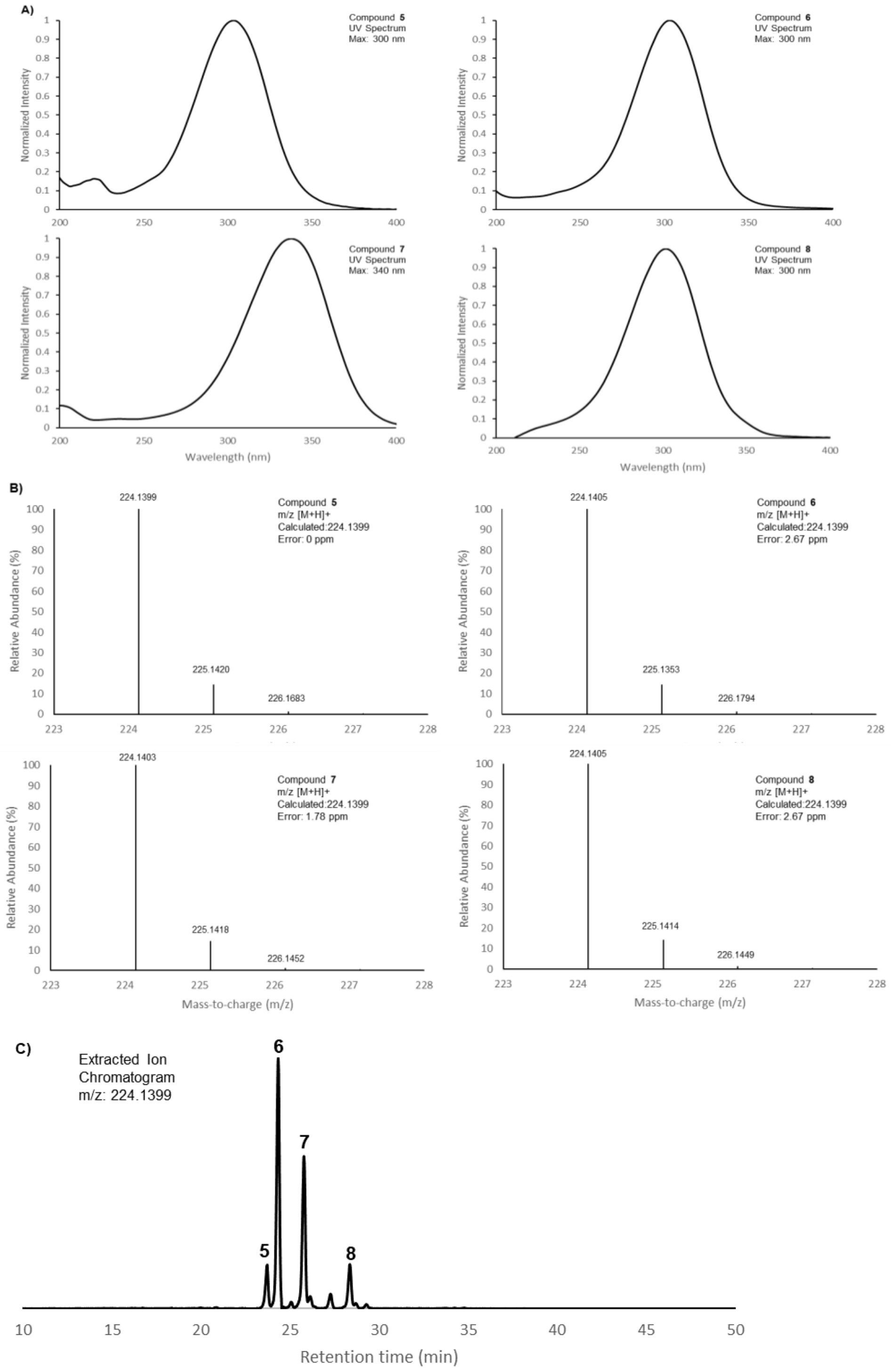


**Figure S7.** Production of triacsin derivatives by *S. tsukubaensis*. A) UV spectra for **5-8**. B) HRMS for **5-8.** C) EIC showing production of **5-8** by *S. tsukubaensis.*

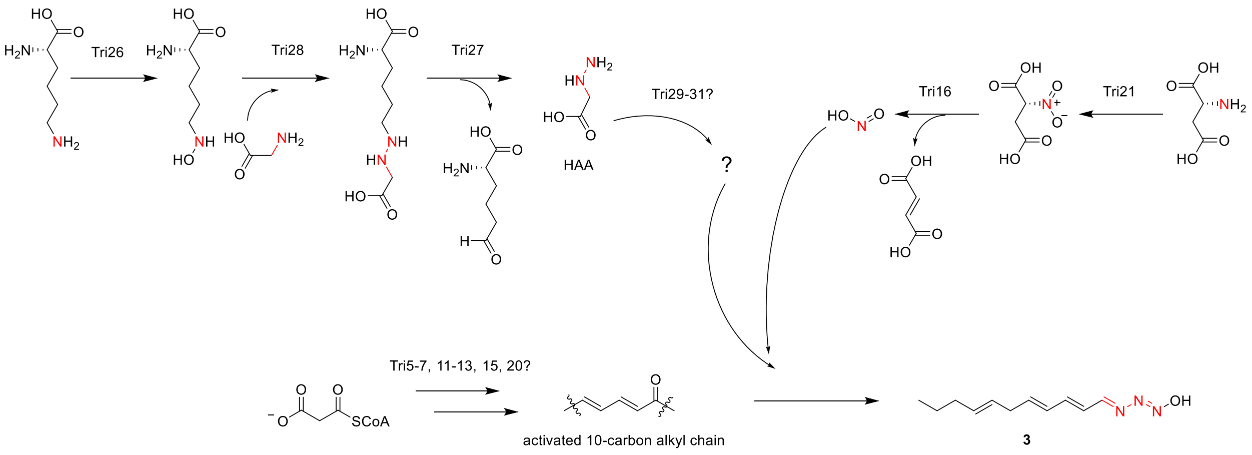


**Figure S8.** Proposed biosynthetic pathway for triacsin C.

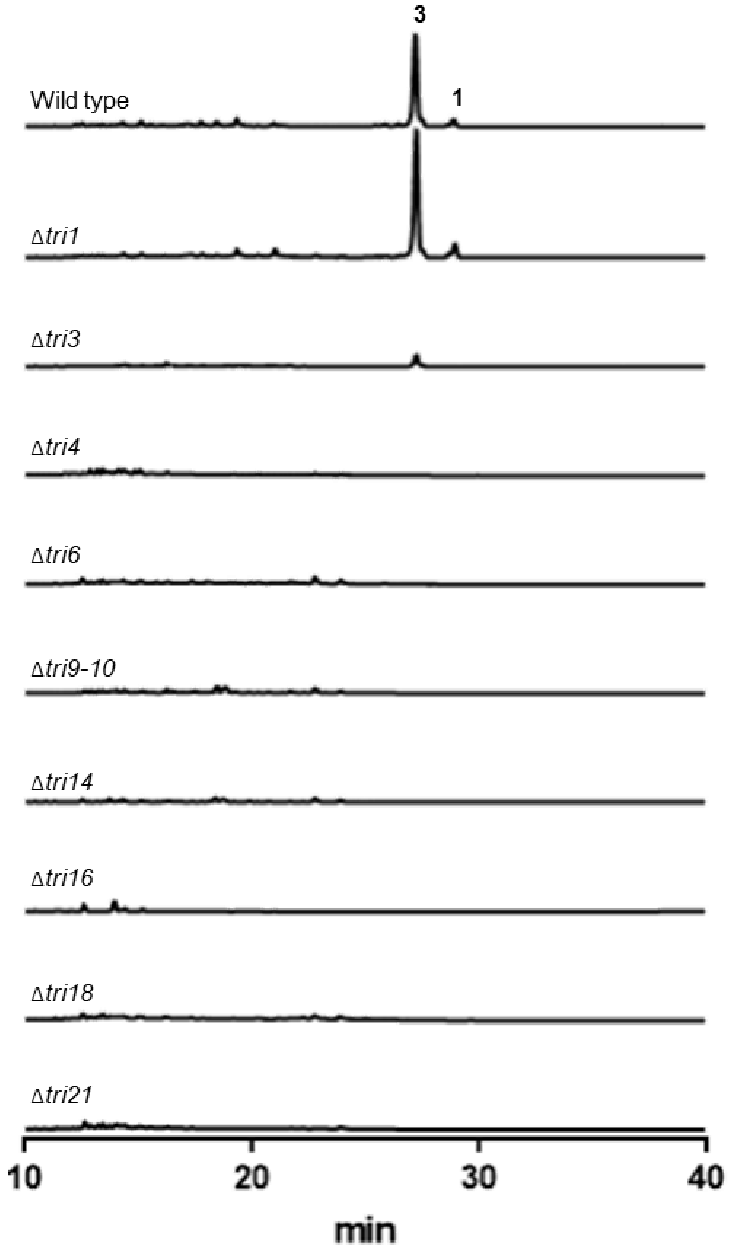


**Figure S9.** HPLC-UV analysis (300 nm) of all gene disruptions in *S. aureofaciens.*

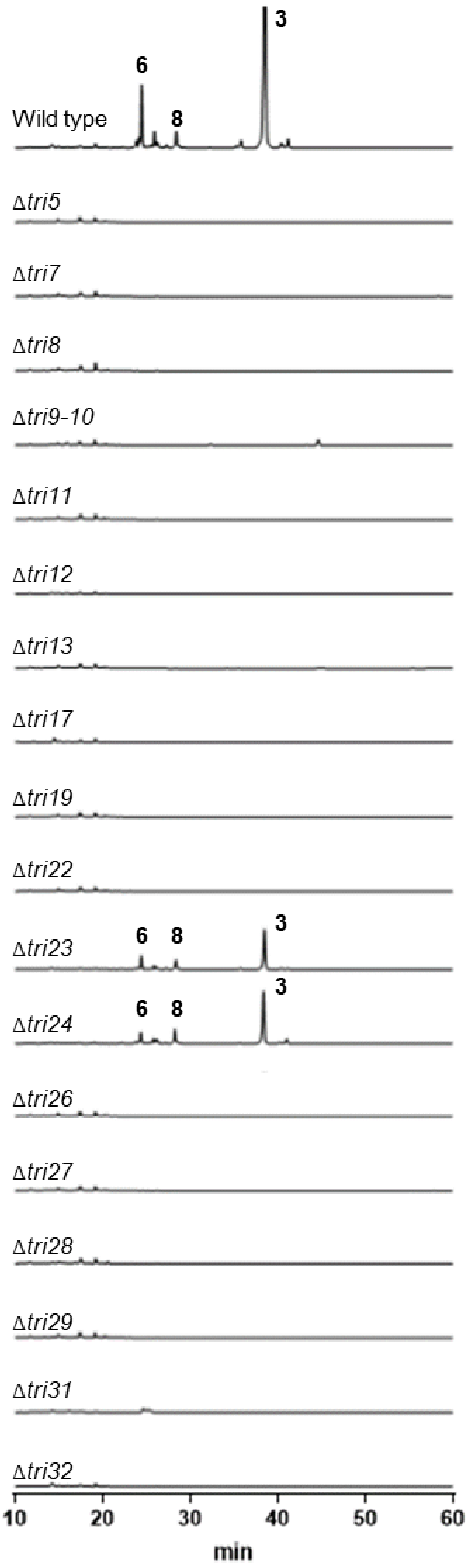


**Figure S10.** HPLC-UV analysis (300 nm) of all gene disruptions in *S. tsukubaensis.*

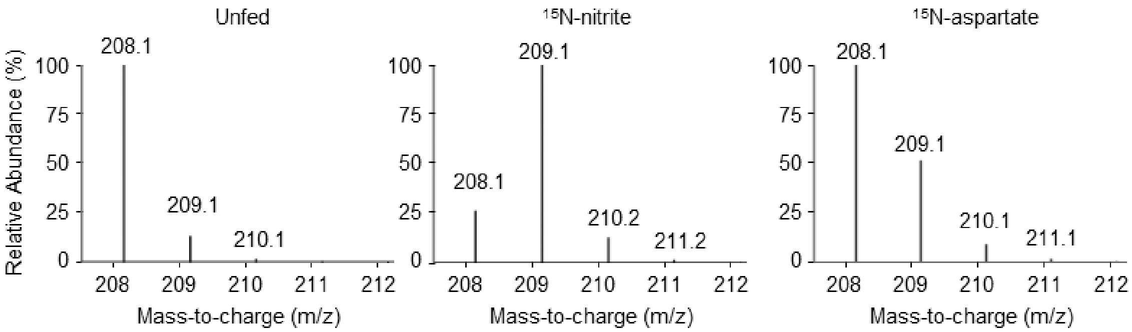


**Figure S11.** LC-MS isotopic peak analysis of **3** after $^{15}$N-nitrite and $^{15}$N-aspartate precursor feeding studies. Wild type cultures of *S. aureofaciens* were supplied with 1 mM $^{15}$N-nitrite or 10 mM $^{15}$N-aspartate and prepared as described in the experimental section.

A)

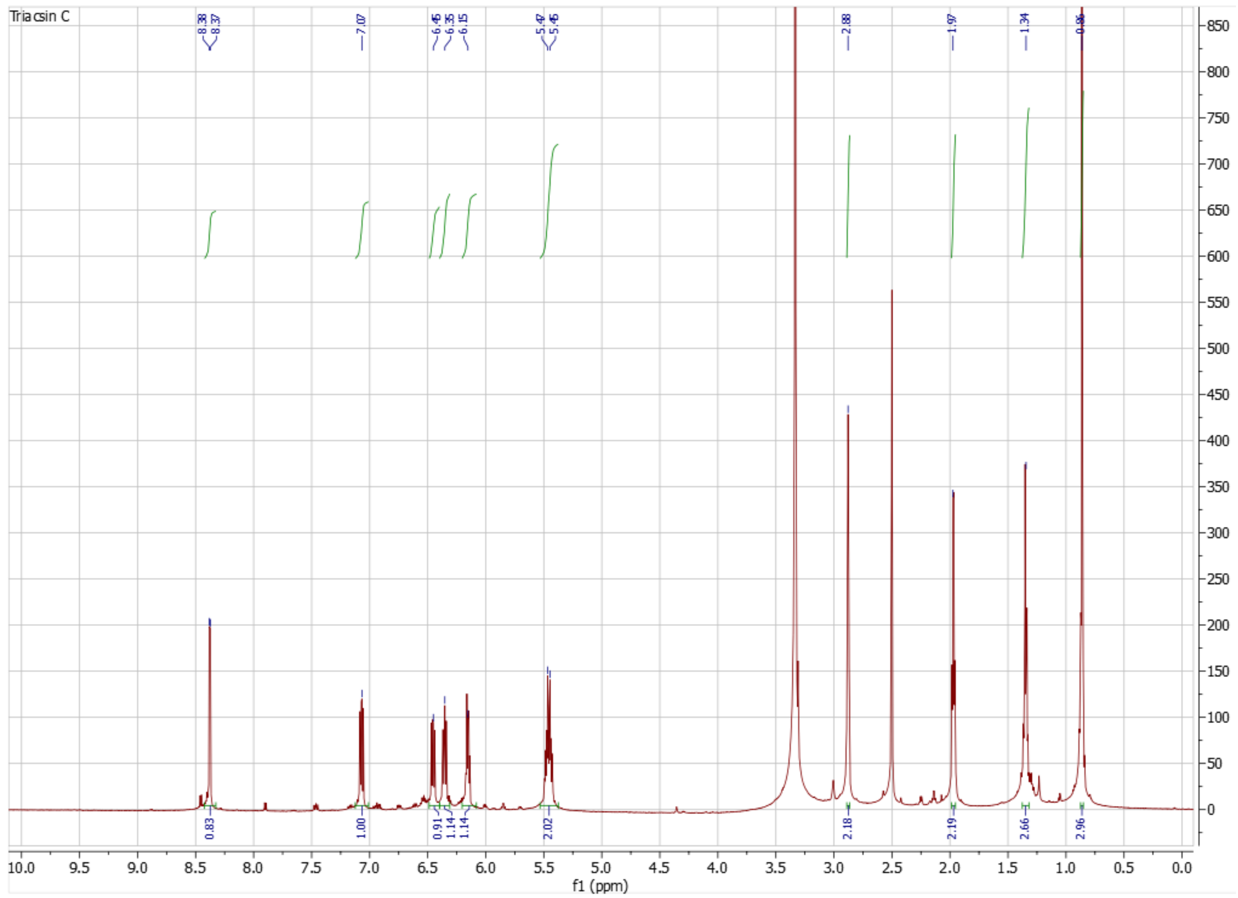


**Figure S12.** NMR characterization of **3** purified from cultures fed unlabeled glycine**.** A) $^{1}$H NMR spectrum (DMSO-$d_6$, 900MHz) of **3**.

**B)**

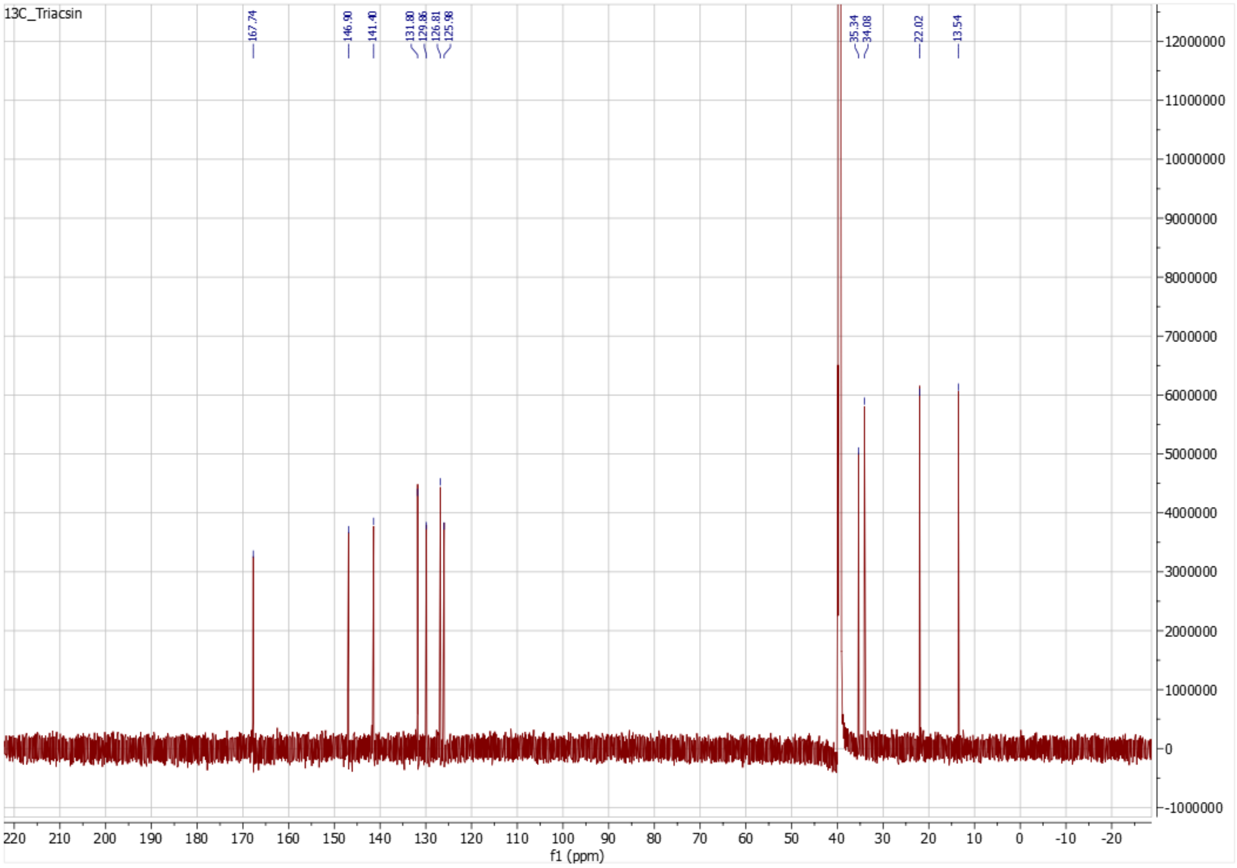


**Figure S12.** B) $^{13}$C NMR spectrum (DMSO-$d_6$, 226MHz) of **3**.

C)

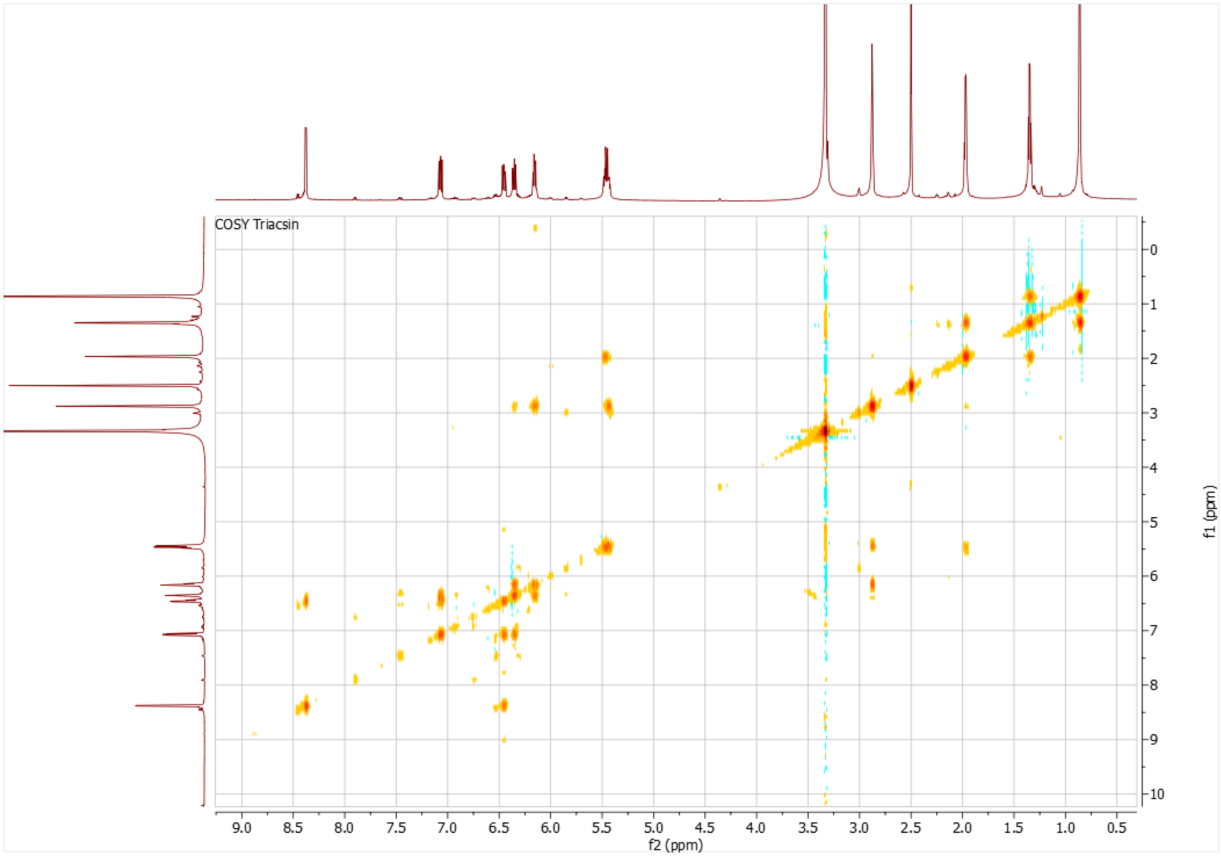


**Figure S12.** C) $^1$H, $^1$H COSY spectrum (DMSO-$d_6$, 900MHz) of **3**.

D)

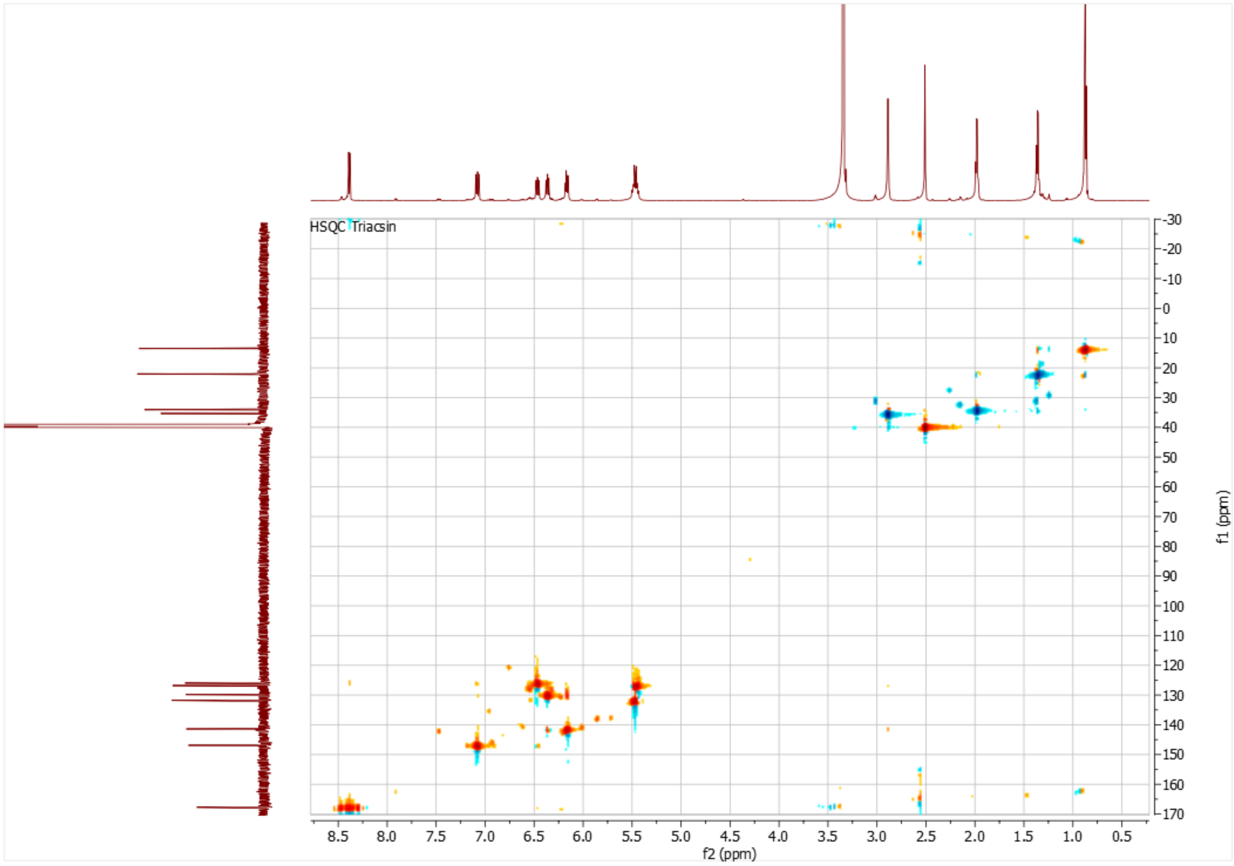


**Figure S12.** D) $^{1}H$, $^{13}C$ HSQC spectrum (DMSO-$d_6$, 900MHz) of **3**.

E)

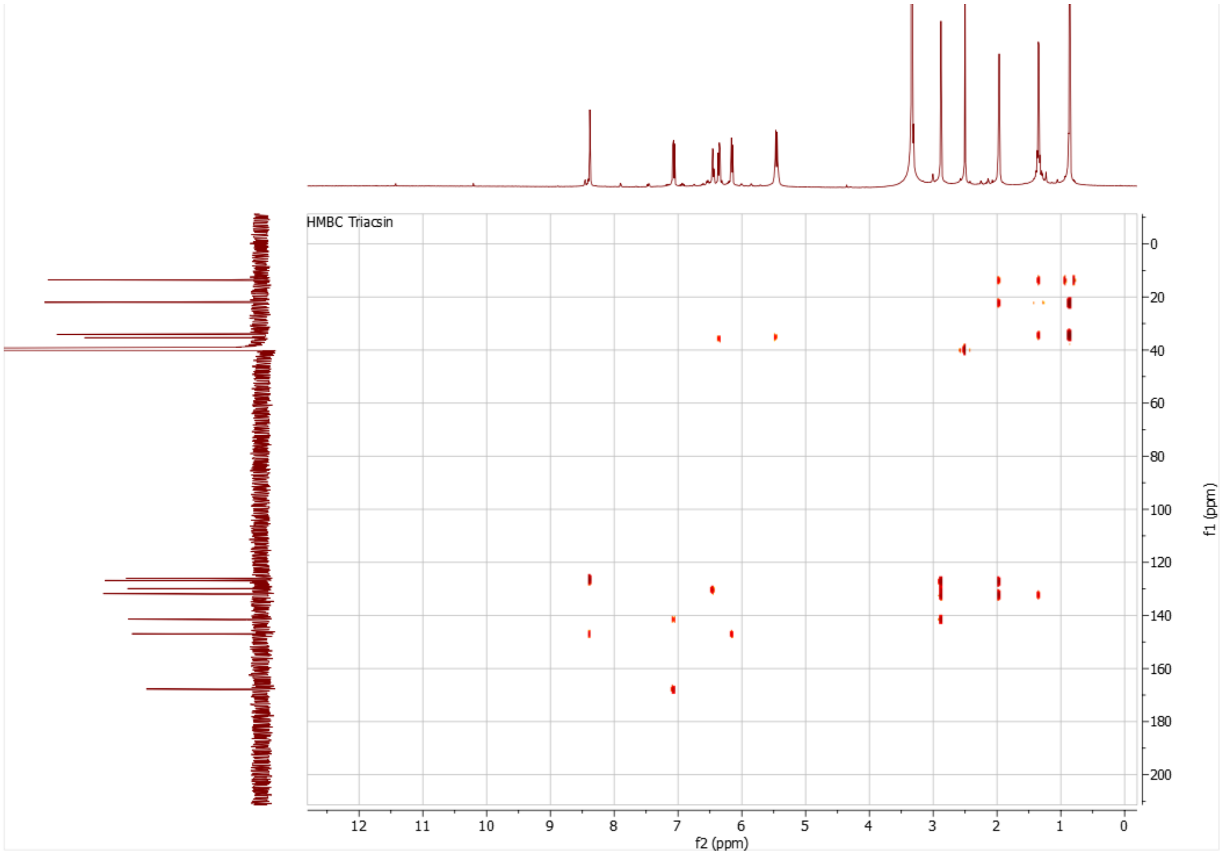


**Figure S12.** E) $^{1}H$, $^{13}C$ HMBC spectrum (DMSO-$d_6$, 900MHz) of **3**.

A)

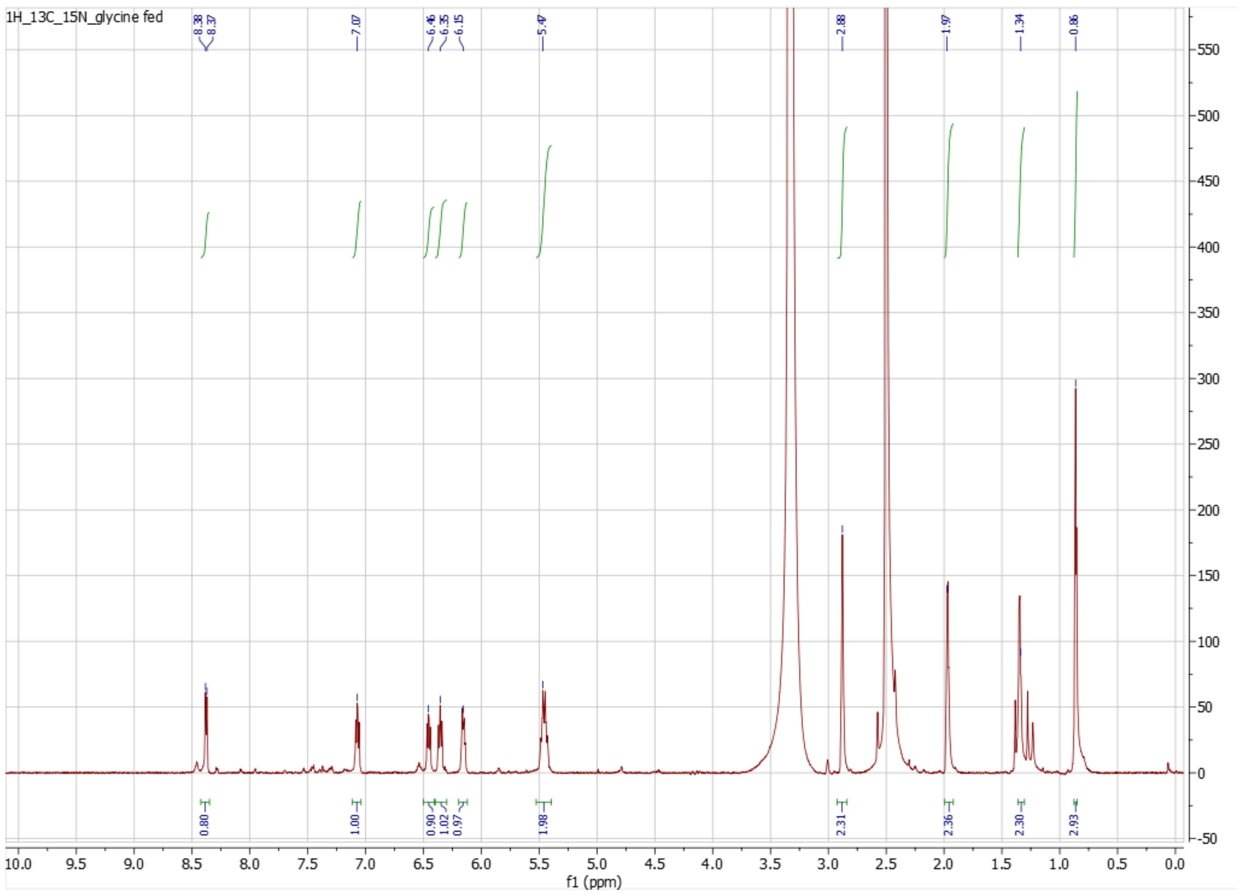


**Figure S13.** NMR characterization of **3** purified from cultures fed 2-$^{13}$C, $^{15}$N-glycine. A) $^{1}$H NMR spectrum (DMSO-$d_6$, 900MHz) of labeled **3**.

**B)**

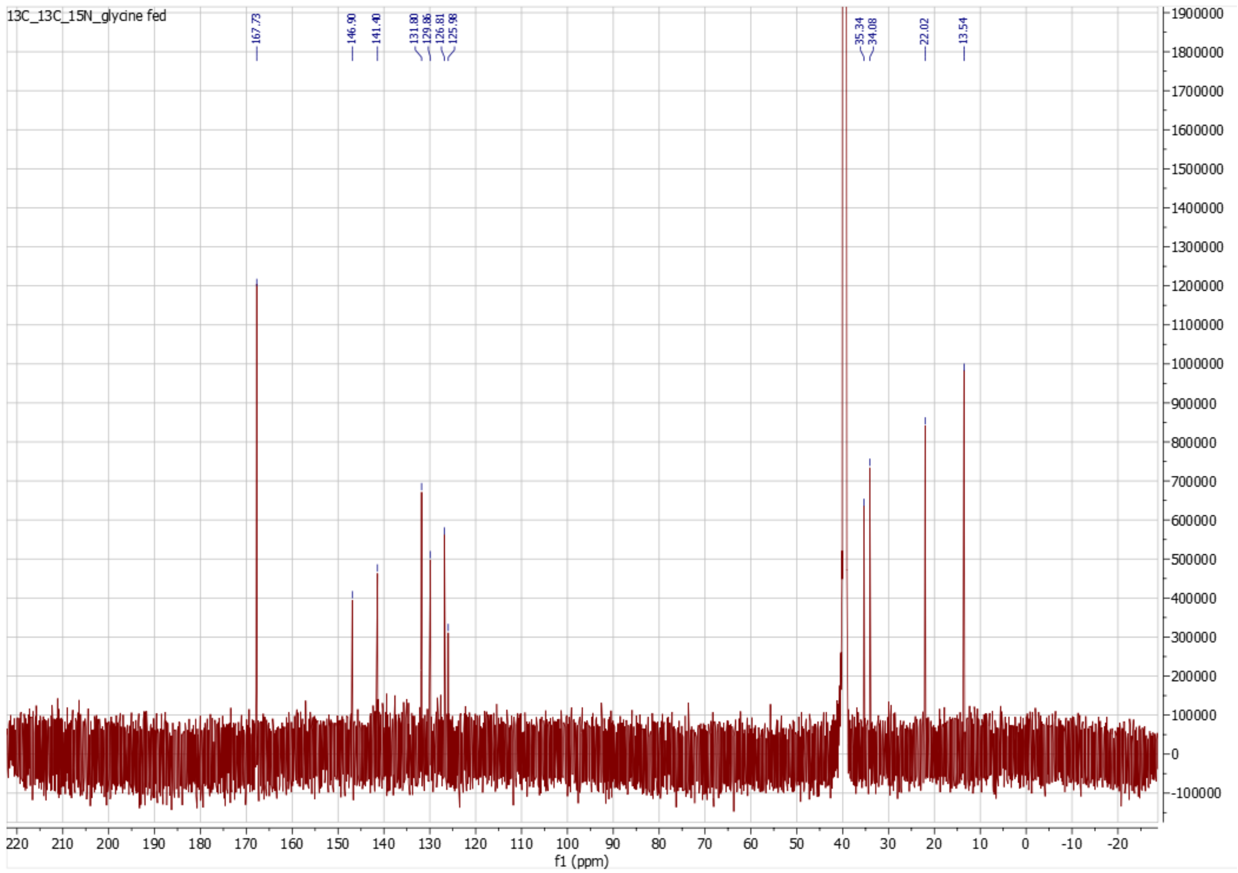


**Figure S13.** B) $^{13}C$ NMR spectrum (DMSO-$d_6$, 226MHz) of labeled **3**.
